# Is variation in female aggressiveness across *Drosophila* species associated with reproductive potential?

**DOI:** 10.1101/2024.08.27.609931

**Authors:** Eleanor Bath, Jennifer M. Gleason

**Affiliations:** Lady Margaret Hall, University of Oxford, Oxford, United Kingdom; Ecology and Evolutionary Biology, University of Kansas, Lawrence, Kansas, USA

**Keywords:** female aggression, Drosophila, sexual selection, social selection, machine learning

## Abstract

Aggression is a key determinant of fitness in many species, mediating access to mates, food, and breeding sites. Variation in intrasexual aggression across species is likely driven by variation in resource availability and distribution. While males primarily compete over access to mates, females are likely to compete over resources to maximize offspring quantity and/or quality, such as food or breeding sites. To date, however, most studies have focused on male aggression and we know little about drivers of female aggression across species. To investigate potential reproductive drivers of female aggression, we tested the relationship between three reproductive traits and aggression in eight Drosophila species. Using machine learning classifiers developed for *D. melanogaster*, we quantified aggressive behaviours displayed in the presence of yeast for mated and unmated females. We found that female aggression was correlated with ovariole number across species, suggesting that females that lay more eggs are more aggressive. A need for resources for egg production or oviposition sites may therefore be drivers of female aggression, though other potential hypotheses are discussed.

## Background

Same species aggression is almost ubiquitous throughout the animal kingdom. Aggression can be a key determinant of fitness, establishing an individual’s social dominance, access to resources (mates, food, shelter, and/or territory), and vulnerability to predation [1]. Intrasexual aggression has been extensively studied in males but has been relatively neglected in females [2], despite evidence that females of a wide range of taxa compete aggressively over a diversity of resources [e.g. food, nesting sites, territory, 1, 3, 4] with consequences for survival and reproduction [1, 4, 5]. Individuals are predicted to be most motivated to fight (and continue fighting) when they perceive value in a contested resource [6]. The value of a resource differs between individuals, and within an individual’s lifetime. The reproductive stage of a female modifies how valuable resources are at any given time, influencing her likelihood of engaging, persisting in, and escalating aggressive contests [7–9]. For some species, resources required for offspring production and provisioning (e.g. food, nesting sites) may be the most limiting resource, causing aggression to peak at times when fecundity is at a maximum [9–11]. For other species, mates or the resources they provide (e.g. sperm, nuptial gifts, paternal care) may be a limited resource, resulting in increased female competition before mating for access to mates [12–14].

Although within-species studies demonstrate that females fight predominantly over resources for reproduction, we have few testable hypotheses about the relationship between reproductive resources and patterns of female aggression among species. The only comparative study (to our knowledge) found that cavity-nesting bird species are more aggressive than open-nesting sister species, suggesting selection for females to fight over limited nesting sites [15]. By framing resources in terms of their value to females, we can predict when females should compete both within and across species, allowing us to systematically test potential drivers of female aggression. We therefore hypothesize that the mating system and reproductive potential of a species may explain patterns of female aggression, because they shape how females value contested resources such as food. We tested this hypothesis with eight species of *Drosophila* that vary in reproductive potential and mating systems in fights over a same food resource.

### Drosophila aggression

In *Drosophila melanogaster,* mated females fight more than unmated females over food [16]. The change in aggression after mating is stimulated by the transfer of sperm and seminal fluid proteins during mating [16] and may be driven by an increased need for protein-rich food to fuel egg production in mated females [17]. Female aggression also evolves in response to the social environment, suggesting increasing aggression is an adaptive response to increasing competition [18]. The clear relationship between reproduction and aggression in *D. melanogaster* sets the groundwork to examine the relationship between variation in reproductive traits and aggression across species with different reproductive strategies.

The genus *Drosophila* encompasses over 2000 species, which vary in their mating behaviour [19–22]. Female fecundity is determined by the number of ovarioles (egg-producing compartments in the ovaries), each of which has the capacity to produce two eggs per day [23]; species with more ovarioles produce more offspring than females of species with few ovarioles [24]. Because females need protein to produce eggs after mating [25], competition for protein-rich food may lead to the possibility that variation in reproductive potential (i.e. ovariole number) across species is associated with aggressive responses [22] due to increased need or valuation of the contested resource.

Females of some *Drosophila* species mate multiply each day whereas females of other species have a refractory period or may never remate [19]. The quantity of sperm transferred affects female propensity to remate; females who receive few sperm need to remate often to maximize their reproductive potential [26]. Across species, the amount of sperm transferred is inversely related to the size of sperm [27–29]. Sperm investment, either in size or number of sperm, varies across the genus [27]. Variation in remating rate and sperm provisioning might result in differences among species in the relationship between a female’s mating status and her investment in aggression.

In this study, we tested if reproductive traits are related to female aggression in *Drosophila*. Aggression is understudied in non-melanogaster *Drosophila* species, with a few exceptions [30–34]. Most *Drosophila* studies have focused on male aggression, though unmated females have been examined in 10 species other than *D. melanogaster* [31, 35]. We used a comparative approach among *Drosophila* species to test the predicted drivers of female aggression across species: ovariole number, sperm length and remating rate. We chose *Drosophila* species spanning the genus with variation in both ovariole number and sperm size.

We focused on aggression over a food (yeast) resource because this form of competition is common across taxa [4]. The more females invest in reproduction, the greater their energetic and time demands, leading to increased consumption of food, as well as increased intake of specific nutrients (e.g. protein, specific amino acids [36, 37]). Because females need food for investment in reproduction, females should be motivated to fight over access to food when it is limited, leading to a positive correlation between investment in reproduction and aggression over food. Thus, as the demands of reproduction increase, females have increased resource needs and the resource has increased value. When females compete for food, females should be most aggressive when energetic demands are highest. We can use this model to predict patterns of aggression across species: species that have high reproductive energy demands (i.e. produce offspring at high rates) will be more aggressive over food than species with low demands.

Resource valuation leads to several predictions (figure 1): 1) Within species, mated females will show more aggression than unmated females because mated females produce eggs and need oviposition sites. Producing eggs requires consumption of food, driving increased aggression after mating in all species (figure 1A); 2). Among species, species with more ovarioles will display more aggression than those with fewer (figure 1B); greater egg production increases resource requirements; 3) Across species, we expect a negative correlation between mated female aggression and sperm length (figure 1C) and mated female aggression. Species with longer sperm transfer fewer sperm in each mating, meaning that fewer eggs are fertilized than in species with shorter sperm. We would therefore expect females receiving few, long sperm to have low energetic demands for egg laying and therefore show lower rates of aggression than females of species who receive many, short sperm. 4). Because remating rate is positively correlated with sperm length, we expect to see a negative correlation between remating rate and female aggression and change in aggression after mating (figure 1D). 5) Among species, the ratio of mated to unmated female aggression will be greater in species with higher resource demands than in species with low resource demands, thus the change in aggression with mating will be positively correlated with ovariole number (figure 1E) and negatively correlated with sperm length and aggression rate (figure 1E-G).

**Figure 1:**
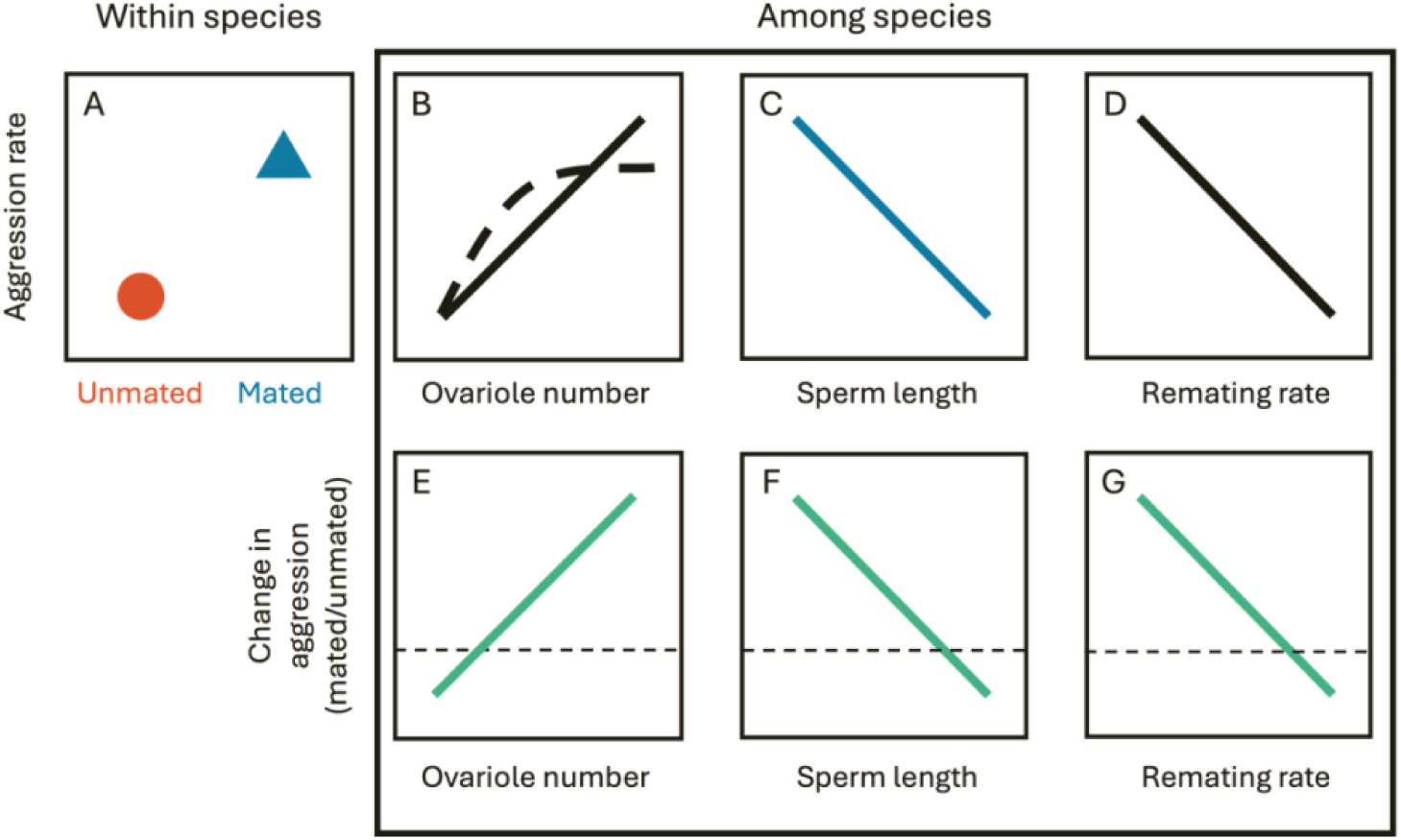
Predictions for the relationship between aggression and potential drivers of aggression across species. See text for detailed arguments. In B, **t**he exact nature of the relationship between ovariole number and aggression rate is plotted as a solid line to indicate a linear relationship and a dashed line to indicate a situation where a maximum level of aggression is reached and cannot increase any further. For E-G the dashed line indicates equal aggression in mated and unmated females. Above the line indicates mated females are more aggressive than unmated females.

To measure female aggression in *Drosophila* species, we used machine learning to automatically score headbutting and fencing in mated and unmated females. The development of software to track animal locomotion and behaviour has improved our ability to study broad-scale and individual-level behaviours [38], but the use of such analyses identifying and classifying behaviours is usually focused on one species at a time [though see 39, 40]. Here we examined eight *Drosophila* species with a range of phylogenetic relatedness, and differences in reproductive traits, to investigate cross-species relationships with reproductive drivers of aggression.

## Methods

### Strain maintenance and unmated fly collection

All strains (table S1) were maintained on standard cornmeal-molasses food in multiple 75 ml bottles at 25°C on a 12:12 light:dark cycle. Unmated males and females were collected on ice within a day of eclosion except for species that are sexually mature in a day or less (*D. willistoni* and *D. nebulosa*), which were collected within 7 hours of eclosion. Males were housed in 12.5 ml vials in groups of 10 with *ad libitum* access to food. Females were housed individually.

### Aggression assay

The aggression assay followed standard methods [16]. We performed all experiments after sexual maturity (table S1). To compare unmated and mated female aggression, the day before the aggression assay, we paired a random subset of the unmated females with one or two males of their species at the start of the light cycle until the female mated or two hours elapsed. Any female that mated was transferred alone to a fresh food vial to be kept overnight. Females that did not mate were discarded. Unmated females that were never exposed to males were moved to fresh food vials at the same time as mated females to be kept in identical conditions overnight.

At 24-hrs after mating, females were starved for two hours to increase their motivation to compete for food [41]: each mated and unmated female of a single species was placed in an individual empty vial with a moist cotton ball. Each female was then paired with a female from the same treatment (mated or unmated) in a contest arena (20 mm in diameter), with apple juice agar and a small patch of yeast paste (Supplementary Video). Up to 20 dyads of a single species were video recorded simultaneously with a Basler ac A2440-75µm camera. Where possible, equal numbers of mated and unmated dyads were included in each video to control for time-of-day effects. Pairs were given 5 minutes to acclimatize to the arena. We recorded videos for 15 minutes using Kinovea software (v 0.9.5; https://www.kinovea.org) at 30 frames-per-second.

### Automated tracking and machine learning methods

To score behaviours, we used automated tracking and machine learning with each video [42]. Briefly, the Caltech fly tracker [43] determined the location and per-frame characteristics (e.g. velocity, distance to other flies, orientation) of each fly within an arena. Each species was tracked separately, with species-specific tracking parameters (e.g. background and foreground thresholds) to account for differences in body size. Videos were inspected manually after tracking to check the accuracy of tracking. Videos that did not track correctly were retracked with video-specific parameters to ensure the highest quality video tracking.

The tracking data were transferred to JAABA [44], where we used supervised machine learning to train classifiers for female aggression. We scored female aggression as headbutts/shoves (attacks using the head and/or body) and fencing (striking the opponent with legs; Supplementary Video). Headbutting is the most commonly used measure of female aggression in *D. melanogaster* studies [16, 45, 46], so we focus our results section on this behaviour, with fencing results reported in the Supplementary Information. Both headbutting and fencing occur in females of multiple species [31, 35], suggesting consistent behavioural repertoires across species. *Drosophila nebulosa* had a wing flicking behaviour that has been observed in courting pairs [47, 48] but because this was only found in one species, we have not included analyses here.

We used an existing classifier for headbutts previously trained on *Drosophila melanogaster* [42]. We trained a new classifier for fencing and a new locomotion classifier, defined as any time a female moved around the arena. All classifiers were ground-truthed by manually scoring a subset of videos with the scorer blind to the automated classifier score. Ground-truthing was conducted on each species to test for differences in accuracy among species. We used 30 randomly chosen intervals for each behaviour in each species, using the ‘balanced random’ option in JAABA, which ensures that some of the intervals contain the behaviour. We selected bout lengths (the number of frames an instance of that behaviour was scored) the same length as the average bout length for each species, because bout lengths differed among species (table S2). As with most behaviours, some instances of behaviours were clear (‘Certain’), while others were more difficult to classify (‘Difficult’). When manually scoring during ground-truthing, we scored the frames as either containing the behaviour (‘Behaviour’) or not (‘None’), and within each of those two groups, whether the behaviour was easy to distinguish (‘Certain’) or difficult to score (‘Difficult’), resulting in four possible outcomes for a frame: ‘Behaviour Certain’, ‘Behaviour Difficult’, ‘None Certain’, ‘None Difficult’. We separately analysed the frames where the behaviour was clear from those where it was difficult, because we expected differences in the performance of the classifier between ‘Certain’ and ‘Difficult’ cases.

To quantify behaviours, we obtained the number of frames in which JAABA scored a focal individual in a dyad displayed that behaviour. We divided this number of frames by the total number of frames in which the individual was tracked to produce the proportion of frames an individual engaged in the behaviour. Using the proportion of frames allowed us to account for slight differences in frame numbers among videos or individuals, due to recording or tracking discrepancies. We calculated values for each individual in a dyad, but for some analyses (see below), we calculated a combined score for the dyad based on both individuals’ tracking data.

For all behavioural classifiers, we used the proportion of frames scored for a behaviour as our response variable (e.g. proportion of frames scored as headbutting). The proportion of frames in which one individual displayed aggression (headbutting and fencing) was positively correlated with the proportion of frames her opponent displayed aggression (figures S1 & S2), thus counting all frames with aggression led to pseudoreplication. To avoid pseudoreplication, we summed the number of frames for both individuals in a dyad to calculate the total number of frames in which the behaviour was observed in the arena to produce one value per dyad. We divided this number by the number of frames in the video to produce the total proportion of frames in which the behaviour happened for the pair. We then used this composite value for each pair, resulting in only one value per dyad. For locomotion behaviour, how much one individual moved was correlated with how much her opponent moved (figure S3). However, both individuals could be moving at the same time so combining the number of frames in which each individual moved would have produced an overestimate of locomotion within a pair. We therefore randomly chose one individual from each dyad and ran the analysis on those individuals to avoid pseudoreplication.

### Phylogenetic tree and species data

We built a phylogeny for our eight *Drosophila* species using the Open Tree of Life data store [49] and custom synthesis tools [50], based on phylogenies from multiple sources [24, 51–53]. Branch lengths were estimated from date using Chronosynth [54]. Once a tree containing 200 species had been built for another research project, we trimmed this tree to contain only our eight species (table S1, figure S4). We attempted to account for phylogeny by pairing species, when possible, in the same species group (i.e. *D. ananassae, D. bipectinata* in the melanogaster group, *D. willistoni, D. nebulosa* in the willistoni group, *D. saltans, D. sturtevanti* in the saltans group).

We collected values on ovariole number, sperm length, and remating rate from the literature (table S1). For several of our species, we did not have data for ovariole number or remating rate. To measure remating rate, we mated females as described above. Once those pairs had finished mating, we removed the male and left the female in the vial alone overnight. Twenty-four hours later, we placed two new unmated males in the vial with the female. Flies were observed for two hours to determine if remating occurred [following 55, 56]. Our sample sizes were 29 *D. saltans* vials (4 remated), 37 *D. sturtevanti* (37 remated), and 48 *D. willistoni* vials (1 remated). We attempted to collect these data for *D. nebulosa* but could not get a large enough sample size. We then calculated the percentage of females that remated for use in our analysis of remating rate.

We counted the number of ovarioles in species for which we could not find ovariole number in the literature. We followed the procedures outlined in Lobell et al. [57]. Briefly, unmated females were collected and housed individually. When sexually mature, a female was placed on food with yeast paste overnight before freezing at −20°C. Within a month, females were dissected using fine needles in a drop of PBS and stained in a solution of PBS saturated with crystal violet (Thermo Scientific). Ovarioles were counted from both ovaries. Strains measured are given in table S1. At least 20 females were dissected from each species.

### Data analysis

All analyses were run in R v4.2.1 [58]. To analyse our behavioural data, we used generalized linear models (GLMs) from the *lme4* package [59]. Model fit and assumptions were checked by visual examination of diagnostic plots. We acquired p-values using the ‘Anova’ function from the *car* package [60]. To test if classifier accuracy was correlated with the frequency of the behaviour, we conducted a linear model with classifier accuracy (% of instances correct) as the response variable and the proportion of frames in which the behaviour was observed as the explanatory variable. To analyse the effects of species and mating status, we used the total proportion of frames in which flies performed the behaviour as the response variable, with species, mating status, and their interaction as explanatory variables:

*Proportion of frames performing behaviour ∼ Species * Mating status*

All variables and their interaction were included in all models as these were key to the experimental design and answering the question at hand. We fitted these models with a quasibinomial error distribution to account for underdispersion in our proportion data. We performed this analysis for all species for headbutting, fencing, and locomotion (tables S3, S4).

To assess if locomotion and aggression were positively correlated, we ran the following GLM with a quasibinomial distribution:

*Proportion of frames aggressive ∼ Proportion of frames moved * Species * Mating status*

To test relationships between aggression and the reproductive characters, we performed phylogenetic analyses using the mean values for each behavioural trait (i.e. headbutting and fencing) for each species and mating treatment as the response variable and the mean value for the reproductive trait (i.e. ovariole number, sperm length, remating rate) as the explanatory variable. To test the change in aggression between mated and unmated females, we calculated the ratio of mated to unmated aggression by dividing the mean species value of the mated behaviour by the mean value of the unmated behaviour (e.g. mated headbutt mean/unmated headbutt mean). We ran a phylogenetic generalized least squares (GLS) models with female median lifespan, average ovariole number, sperm length and remating rate per species as explanatory variables in separate models [nlme package: 61, ape package: 62]. For the analyses with ovariole number, we included average body size as a covariate because ovariole number scales with body size across *Drosophila* species [24, 29]. We log-transformed ovariole number, body size, and sperm length before including them as explanatory variables in phylogenetic GLS models. As we ran four models for each of our response variables; we corrected for multiple testing by applying a Bonferroni correction and reducing the threshold for significance (dividing our significance threshold by the number of models: *p=* 0.05/4 = 0.0125).

## Results

### Accuracy of machine learning across species

We used machine learning classifiers [42] to automatically score headbutts, fencing, and locomotion in eight species. We scored classifier accuracy by ground-truthing a subset of frames within each species (figure S5 and table S2). We evaluated frames in which the behaviour was unequivocally happening or clearly not happening as ‘Certain’ instances and frames where the behaviour was less clear as ‘Difficult’ frames. We used these two categories to allow us to separate clear instances from less clear instances where manual observers often struggle to define behaviours. For the headbutting classifier, accuracy ranged from 67-100% in the ‘Certain’ frames and from 63-100% in the ‘Difficult’ frames. Fencing was more variable with 51-100% for ‘Certain’ frames and 44-100% accuracy in the ‘Difficult’ frames. We included locomotion to investigate differences induced by mating, as increased movement is a post-mating response in *D. melanogaster* [63]. Locomotion is less complicated for the classifier to score because it focuses only on the behaviour of one fly, providing us with a non-aggressive behaviour for assessing classifier accuracy. Locomotion was scored with similar accuracy to headbutts and fencing with a range of 73-100% for the ‘Certain’ frames and 60-100% for the ‘Difficult’ frames (table S2). Overall, headbutting and locomotion were more accurately classified than fencing, though this varied among species (e.g. for *D. nebulosa*, all behaviours had high levels of classifier accuracy, but for *D. hydei* headbutting was more accurately classified than fencing).

We tested if the frequency of behaviours influenced classifier accuracy, on the assumption that species that had more instances of behaviours may be more likely to have higher accuracy. We found that classifier accuracy was not related to how frequently the behaviour was observed for headbutting (F_1,5_ = 0.3, *p* = 0.61) or locomotion (F_1,5_ = 1.03, *p* = 0.35). For fencing, accuracy and frequency were slightly negatively associated (F_1,5_ = 6.76, *p* = 0.041): species with more instances of fencing were less accurate, driven primarily by low accuracy in *D. hydei*. Therefore, the differences among species and behaviours in accuracy were not due to how often the behaviours were displayed.

### Effects of species and mating status on aggression

Results for fencing were broadly the same as for headbutting, thus we only present headbutting results; results for fencing are presented in the online supporting information. We found a significant effect of species on the proportion of frames in which pairs were headbutting (χ^2^_7,538_= 49.76, *p* < 0.001; figure 2). *Drosophila pseudoobscura* was the most aggressive species with respect to headbutting (proportion of frames: 0.018 ± 0.0017) and was significantly more aggressive than all species except *D. saltans* (post-hoc Tukey tests, table S3). Mating status did not have a significant effect (χ^2^_1,538_ = 0.35, *p* = 0.55), nor was there an interaction between species and mating status (χ^2^_7, 538_ = 6.82, *p*= 0.45).

**Figure 2:**
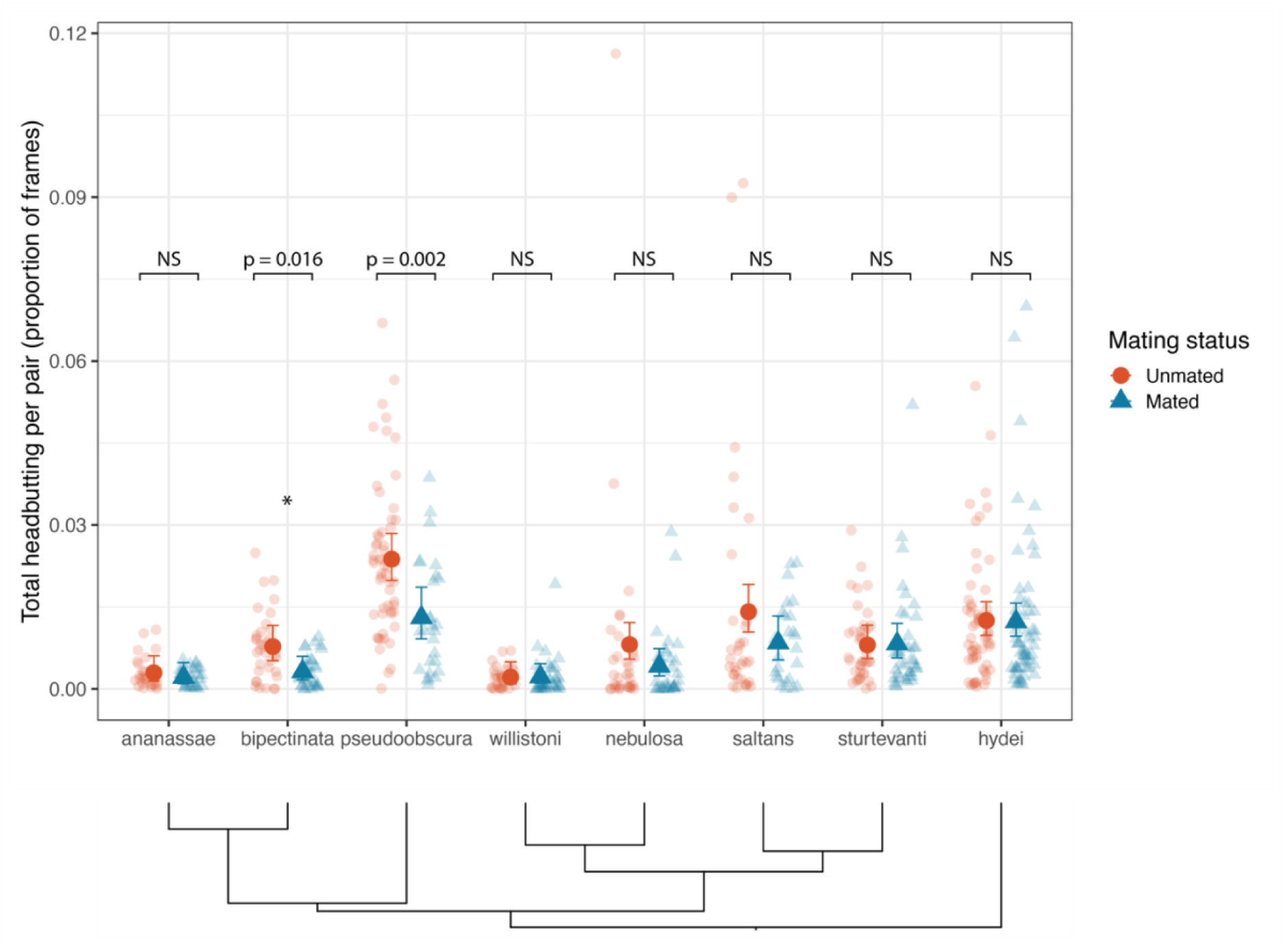
Total proportion of frames labelled with headbutting by species and mating status. Each small red circle represents one dyad of females, with orange circles representing are unmated pairs and blue triangles representing mated pairs. Large points represent model means with error bars representing 95% confidence interval (asymmetric due to proportion data). The phylogenetic relationships among the species are shown underneath the figure (see figure S4 for branch lengths). Sample sizes (unmated, mated): *D. ananassae* (n = 29, 32), *D. bipectinata* (n = 35, 35), *D. pseudoobscura* (n = 57, 27), *D. willistoni* (n = 30, 38), *D. nebulosa* (34, 33), *D. saltans* (n = 34, 25), *D. sturtevanti* (n = 40, 39), *D. hydei* (n = 32, 34).

Although we found no evidence for an interaction between species and mating status, visual inspection of the data revealed trends in how species responded to mating. We performed pairwise comparisons within each species between mated and unmated females. Unmated females were significantly more aggressive than mated females in *D. bipectinata* (Odds Ratio Unmated/Mated = 2.46, z = 2.409, *p* = 0.016; figure 3) and *D. pseudoobscura* (OR = 1.84, z = 3.065, *p* = 0.002). Trending in the same direction, though not significantly, were *D. hydei* (OR = 1.55, z = 1.672, *p* = 0.0945), *D. nebulosa* (OR = 1.96, z = 1.943, *p* = 0.052), and *D. saltans* (OR = 1.68, z = 1.898, *p* = 0.058).

**Figure 3:**
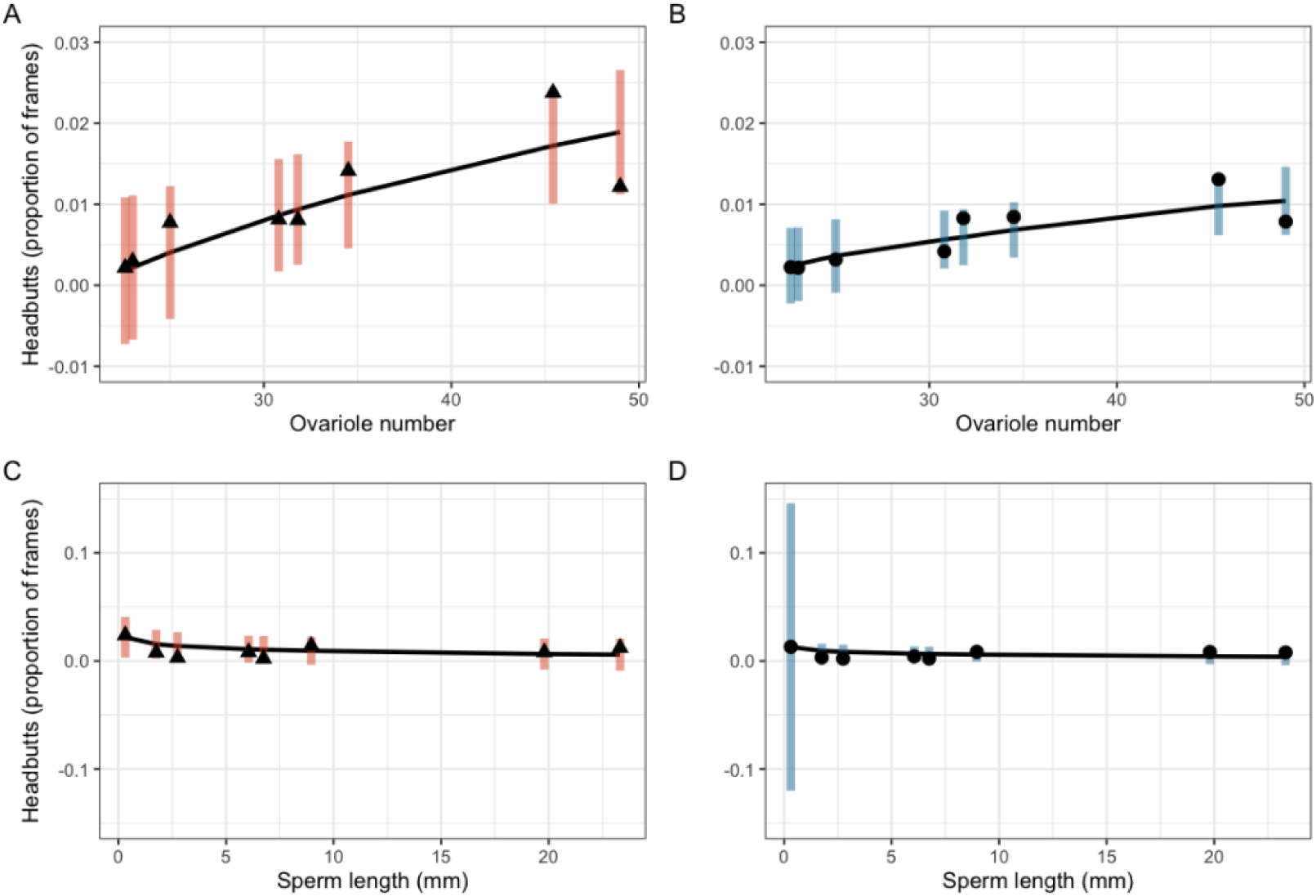
Relationship between headbutts and fecundity traits. Relationship between headbutts and ovariole number in A) unmated females, B) mated females and relationship between headbutts and sperm length in C) unmated females, D) mated females. Dots (triangles in A and C and circles in B and D) represent the species mean proportion of frames in which headbutting was recorded. Confidence intervals represent 95% confidence intervals calculated in the phylogenetic analysis including body size as a covariate. The black line represents the model predictions.

To investigate the effect of phylogeny on headbutting, we ran a phylogenetic GLS model with a phylogeny containing our eight species for mated and unmated headbutts separately. We found that both mated and unmated headbutting had a significant phylogenetic signal with λ = 1.23 and λ = 1.07 respectively. Closely related species were similar in their levels of headbutting for both mated and unmated females. However, the relative change in aggression after mating did not have a significant phylogenetic signal (λ = − 0.24).

### Locomotion

Species and mating status had a significant interaction with respect to the proportion of frames in which individual flies were moving (χ^2^ = 34.569, *p* < 0.0001; figure S6). Unmated females moved significantly more than mated females in *D. bipectinata* (OR V/M = 5.64, z = 3.804, *p* = 0.0001), but mated females moved more than unmated female *D. sturtevanti* (OR = 0.42, z = − 3.071, *p* = 0.0021). No other species exhibited a significant difference in locomotion with respect to mating status (figure S6). We found no significant effect of locomotion on headbutts, nor were there significant interactions between locomotion, mating status or species (figure S7). Observed differences in aggression among species, therefore, were not attributable to differences in locomotion.

### Ovariole number

Ovariole number and body size were not correlated with each other in our sample of species (F_1,6_ = 3.24, *p* = 0.12). The relationship between ovariole number and the proportion of frames with headbutting was positive (Unmated females: χ^2^_1, 5_ = 40.2, *p* < 0.0001, figure 3A; Mated females: χ^2^_1, 5_ = 14.43, *p* = 0.0001; λ= 1.26, figure 3B). Unmated females had a negative relationship between body size and headbutting—larger species spent less time headbutting (χ^2^_1, 5_ = 8.51, *p* = 0.004; λ= 0.82). This relationship was not seen in mated females (χ^2^_1, 5_ = 0.37, *p* = 0.54). Ovariole number did not predict the change in aggression between unmated and mated flies (χ^2^_1, 5_ = 2.6, *p* = 0.11; λ= −0.68), whereas mating-induced changes in aggression were linked to body size: species with small body sizes experienced a larger change in headbutting between mated and unmated females (χ^2^_1, 5_ = 5.88, *p* = 0.015).

### Sperm length

Sperm length was not correlated with unmated female headbutting (χ^2^_1,6_ = 3.07, *p* = 0.08; λ= 1.17; figure 3C), whereas species with larger sperm tended to have less headbutting in mated females (χ^2^_1,6_ = 20.86, *p* < 0.0001; λ= 1.26; figure 3D). The relative change in headbutting behaviour between mated and unmated females was not associated with sperm length across species (sperm length: χ^2^_1, 6_ = 0.47, *p* = 0.49; λ= −0.69).

### Remating rate

Remating rate and sperm length are correlated in our set of species (F = 8.1, *p* = 0.04, λ= − 0.75). However, remating rate was not related to unmated headbutting (χ^2^_1,5_ = 0.05, *p* = 0.83; λ= 1.09), mated headbutting (χ^2^_1, 5_ = 0.29, *p* = 0.59; λ= 1.24), or relative change in headbutting between mated and unmated females (χ^2^_1, 5_ = 0.65, *p* = 0.42; λ= 0.17).

## Discussion

### Change in aggression

Based on the behaviour of *D. melanogaster,* and consistent with the resource valuation model, we predicted that mated females would be more aggressive than unmated females within species (table 1). In contrast, we found that for many of these species, aggression was unchanged between unmated and mated females or was reduced after mating. We did not see changes in aggression post-mating associated with any of the other traits measured here.

**Table 1.** Results with respect to predictions.

| Predictor variable | Comparison | Predicted relationship | Results of study |  |
| --- | --- | --- | --- | --- |
| Response variable: Change in aggression |  |  |  |  |
| Mating | Within species | Mated > Unmated | No consistent effect across species; in no species were Mated > Unmated;<br>*D. bipectinata* and *D. pseudoobscura*: Unmated > Mated |  |
| Ovariole number | Among species | Positive | None |  |
| Sperm length | Among species | Negative | None |  |
| Remating rate | Among species | Negative | None |  |
| Response variable: Aggression level |  |  | Unmated females | Mated females |
| Ovariole number | Among species | Positive | **Positive** | **Positive** |
| Sperm length | Among species | Negative (mated only) | None | **Negative** |
| Remating rate | Among species | Negative | None | None |

Notes: For the results, only statistically significant relationships are given, or “none” is indicated. Bold text indicates that the hypothesis is supported by the results

Independent of the effect of males on female behaviour, the resource valuation model predicts that females of all species experience an increased demand for protein and other nutrients essential for egg production after mating. Species are likely to vary in their nutritional needs: for instance, *D. melanogaster* females can produce mature eggs on a sucrose-only diet, while *D. hydei* females require protein to synthesize essential proteins and develop eggs to maturity [64]. If nutrient requirements do not change substantially after mating, then aggression may not change. Research investigating the nutritional requirements and patterns of egg production across species are needed to understand how the value of resources changes with female reproductive state, particularly to understand the decrease in aggression seen in two species (*D. bipectinata* and *D. pseudoobscura*).

Male ejaculate components, including sperm and the seminal fluid protein sex peptide, increase aggression in *D. melanogaster* females. The involvement of male ejaculate components in stimulating aggression raises the possibility that female aggression post-mating is a by-product of sexual conflict. However, the nature of male ejaculate components is diverse across the clade: for example, *D. melanogaster* has two sex peptide genes, whereas the *ananassae* group (here represented by *D. ananassae* and *D. bipectinata*) has a proliferation of sex peptide genes [65]. We know little about how the diversity of seminal fluid proteins or sperm translates into changes in post-mating behaviour in females, including aggression. Our results indicate that post-mating changes in aggression and locomotion are not universal.

### Aggression levels among species

Ovariole number determines the number of eggs that can be produced in a day and, consequently, lifetime fecundity [24]. As predicted, we found a positive correlation between ovariole number and aggression for both unmated and mated females (table 1). Because egg production requires a large quantity of protein [66], females who have more ovarioles (and therefore produce more eggs) are likely to require more protein than females producing fewer eggs over the same time. The increased need for a resource (protein) may drive increased aggression, helping to explain the pattern we see across species. However, this pattern may also be explained by differential changes in hormones and physiology across species that are associated with differences in ovariole number or egg production. Alternatively, life-history theory suggests that fast-reproducing species (those with more ovarioles) prioritize current reproduction over future reproduction, thus fast reproducing species may be aggressive to increase their current brood’s chance of success [67].

We predicted that species with more ovarioles show a greater change in aggression after mating than those with fewer ovarioles. However, we found no evidence of this in these species, most likely because most species did not change in aggression in response to mating. The absence of a change may indicate that aggression is not driven by energetic requirements of egg production or species may not value yeast as a food resource. *Drosophila s*pecies vary in their ecology [21], which is likely to generate different selection pressures on female aggression with respect to different resources [1]. However, all the species in this study, with the possible exception of *D. saltans* [68], are generalists that eat and lay eggs on rotting fruit [68, 69]. Yeast should therefore be valuable to all species in our study.

Because *D. melanogaster* female aggressive behaviour is affected by sperm [16], a relationship between mated (but not unmated) female aggression was predicted for sperm length (table 1). In *D. melanogaster* the strength of the post-mating aggressive response may depend on the quantity and/or quality of sperm received [18, 70]. If sperm induction of a female’s response is dose dependent, then females receiving few sperm will have less altered aggressive behaviour (either up- or down-regulated) than those who receive more sperm. Sperm length is inversely related to the quantity of sperm transferred [27, 29], thus aggression should decrease with sperm length, as observed. In contrast to our headbutting results, we found a significant effect of sperm length on changes in fencing after mating (Supplementary Information): females of species with long sperm fenced more after mating than before. Headbutting and fencing may be regulated separately and serve different functions in aggressive interactions.

The number of sperm transferred is related to the remating rate [26]. Because sperm size is inversely related to the number of sperm transferred, females that receive large sperm are more likely to remate than females of species with many small sperm, yet we did not detect a relationship between remating rate and aggression. Thus, the effect of sperm on aggression may be more of a direct effect of the first mating rather than an inherent property of mating system, switching aggression on or off in the first mating and remaining constant regardless of the number of subsequent matings. Regardless, the physiological changes that accompany mating, such as upregulation of juvenile hormone and octopamine signalling [71], may influence female aggression in addition to the potential value of a food resource.

### Evolutionary History

Both headbutting and fencing (Supporting Information) had a high phylogenetic signal, suggesting aggression is conserved among closely related species. Behavioural traits are more evolutionarily labile than morphological or physiological traits [72], although numerous behavioural traits have significant phylogenetic signal [reviewed in 73]. Our study consisted of only eight species, which reduces our ability to detect phylogenetic signal, and some of our species were paired within species groups (i.e. *D. ananassae* and *D. bipectinata*, *D. willistoni* and *D. nebulosa*, *D. saltans* and *D. sturtevanti*), which may artificially inflate our measure of phylogenetic signal. Further studies expanding the range of species in which we study both male and female aggression will give us more insight into the selection pressures and constraints shaping aggression.

### Machine learning for comparative behaviour

We used machine learning classifiers originally developed for *D. melanogaster* for eight different species. To our knowledge, this is the first time machine learning has been used for a comparative analysis of social behaviour in arthropods [for an example of non-social behaviour, see 40]. Our range of accuracies was like other classifiers used only for *Drosophila melanogaster* [42, 74, 75], with some of our accuracies exceeding that of same species classifiers. For our species, classifiers were less accurate in some species, suggesting potential caution and ground-truthing must be used in interpreting data from the same classifier on multiple species.

For most species, in clear instances of headbutting, the classifier was more likely to miss instances of aggression than falsely categorise frames with no headbutting (table S2). Our scores of aggression are, therefore, likely underestimates rather than overestimates of female aggression. For frames in which the behaviour was difficult to categorise, the classifier had an even distribution of false negatives and false positives. We ensured high-quality tracking for all species (see Methods), thus differences in tracking accuracy among species are unlikely to be responsible for differences in classifier accuracy. Differences in classifier accuracy may be due to species-specific differences in headbutting and fencing behaviours. Slight differences in speed, direction, and duration of flies’ movements during aggression may alter the classifier’s ability to score behaviours [42]. Detailed studies are needed for an in-depth analysis of qualitative differences among species in these shared behaviours.

Using the same classifiers on all species allowed us to quickly calculate aggression scores and directly compare species in shared experimental conditions. The ability to use one algorithm for multiple species (with caveats) greatly increases the number of individuals and species that can be scored for complex and subtle behaviours. We therefore demonstrate the potential use of machine learning for comparative behavioural analysis, though caution that a human assessment is needed for ground-truthing and identification of additional behaviours exist that are not shared across species.

## Conclusion

*Drosophila s*pecies vary in female aggression when fighting over a food (yeast) resource. The effect of mating was not universal in these species meaning that post-mating responses need to be examined across the genus. Although our results for the relationship between ovariole number and sperm size with aggression is consistent with a resource valuation model for food, the lack of a change in aggression, or a decrease in aggression post-mating, is not supported by the model. We only tested the model with a single resource, food, and other resources such as mates and oviposition sites may differentially induce aggression according to the mating system and ecology. Differences in physiology associated particularly with ovarioles may be a driver of aggression, but this needs to be tested further. Studies are needed to understand how female behaviour varies and if variation is dependent on ecology, mating system, phylogenetic history, or other aspects of female biology.

## Acknowledgements

This work was supported by NSF IOS-2121849 to JMG. Krish Sanghvi and Biliana Todorova helped with the *Drosophila sturtevanti* mating and remating days. We thank Emily Jane McTavish for help with the phylogeny, Connor McCready for counting ovarioles, and Jonathan Green for comments on the manuscript.

