## Supplementary Material for "Is variation in female aggressiveness across *Drosophila* species associated with reproductive potential?"

Electronic Supplemental Material for:

**Table of Contents:**


Supplementary Material (Tables S1-S4; Figures S1-S8) pages 2-5
Supporting Information (Results for Fencing) pages 16-19

### Supplementary Material

#### Table S1. Species and strains used in the study

| Species | Sub-genus[1] | Species group | Strain[2] | Age of sexual maturity (days) | | Age range (days)[3] | Proportion successful matings (N) | Ovariole number (both ovaries)[5] | Sperm length (mm)[4] | Remating rate (% females that remate in 24 hours)[5] | Lifespan (days) |
|---|---|---|---|---|---|---|---|---|---|---|---|
| | | | | F | M | | | | | | |
| *D. ananassae* | S | melanogaster | 14024-0371.13 | | 4[REF 1] | 5-8 | 0.585 (164) | 23 [REF 2] | 2.75 [REF 3] | 4.23 [REF 1,4,5] | 43 [REF 1] |
| *D. bipectinata* | S | melanogaster | REF 6 | | 2[REF 1] | 5-8 | 0.853 (95) | 20.5[REF 2,7] | 1.75 [REF 1] | 5.46 [REF 1] | 36 [REF 1] |
| *D. hydei* | D | repleta | 15085-1641.68 | 3[REF 8] | 9[REF 8] | 12-19 | 0.965 (173) | 48.92[REF 2,9] | 23.32 [REF 8] | 93.16 [REF 4,10,11] | 55 [REF 12] |
| *D. nebulosa* | S | willistoni | 14030-0761.0 | | 0.5[REF 1] | 4-9 | 0.474 (152) | 30.79[REF 13] | 6.05 [REF 1] | No data | 42 [REF 1] |
| *D. pseudoobscura* | S | obscura | 14011-0121.4 | 3-6[REF 9,14] | 1-5[REF 9,14] | 6-9 | 0.462 (143) | 42.17[REF 2,15, 16] | 0.313 [REF 9,14] | 7.4 [REF 4,10,11,14,16] | 54 [REF 17] |
| *D. saltans* | S | saltans | 14045-0911.00 | | 3[REF1] | 6-9 | 0.6 (145) | 34.5[REF 13] | 8.96 [REF 1] | 13.79 [REF 13] | 59 [REF 1] |
| *D. sturtevanti* | S | saltans | Multiple lines[6] | | 9[REF 1] | 10-15 | 1 (119) | 31.81[REF 13] | 19.8 [REF 1] | 100 [REF 13] | 36 [REF 1] |
| *D. willistoni* | S | willistoni | L'Habitatué[REF 18] | | 0.5[REF 1] | 4-7 | 0.898 (157) | 22.6[REF 2] | 6.745 [REF 1] | 2.08 [REF 13] | 28 [REF 1] |

[1]Subgenus: S= Sophophora, D= Drosophila
[2]Numbers are the National Drosophila Species Stock Center (NDSCC) strain IDs.
[3]Days since eclosion when the experiments were conducted.
[4]*D. pseudoobscura* has 2-3 types of sperm that are different sizes. For our analyses, we used the longest sperm length as these have been shown to be the fertilising sperm type.
[5]When there are multiple references, the values are the means of multiple studies.
[6]The *D. sturtevanti* line used was a mass bred population started from round robin mating of five NDSSC lines: 14043-0871.01, 14043-0871.05, 14043-0871.07, 14043-0871.15 and 14043-0871.MX9-36

*References:*

10. Markow T.A. 2002 Perspective: Female remating, operational sex ratio, and the arena of sexual selection in *Drosophila* species. *Evolution* **56**(9), 1725-1734.
11. Markow T.A., Beall S., Castrezana S.J. 2012 The wild side of life Drosophila reproduction in nature. *Fly* **6**(2), 98-101. (doi:10.4161/fly.19552).
12. Promislow D.E.L., Haselkorn T.S. 2002 Age-specific metabolic rates and mortality rates in the genus *Drosophila*. *Aging Cell* **1**, 66-74.
13. This paper
14. Holman L., Freckleton R.P., Snook R.R. 2008 What use is an infertile sperm? A comparative study of sperm-heteromorphic *Drosophila*. *Evolution* **62**(2), 374-385. (doi:10.1111/j.1558-5646.2007.00280.x).
15. Moreteau B., Gibert P., Pétavy G., Moreteau J.-C., Huey R.B., David J.R. 2003 Morphometrical evolution in a Drosophila clade: the *Drosophila obscura* group. *J Zool Syst Evol Res* **41**, 64-71.
16. Snook R.R., Markow T.A. 2001 Mating system evolution in sperm-heteromorphic *Drosophila*. *J Insect Phys* **47**(9), 957-964. (doi:10.1016/S0022-1910(01)00070-1).
17. Yoon J.S., Gagen K.P., Zhu D.L. 1990 Longevity of 68 species of *Drosophila*. *Ohio J Sci* **90**(1), 16-32.
18. Gleason J., Griffith E., Powell J. 1998 A molecular phylogeny of the *Drosophila willistoni* group: Conflicts between species concepts? *Evolution* **52**(4), 1093-1103. (doi:10.2307/2411239).

**Table S2: Ground-truthing results all behaviour classifiers by species**

We classified three behaviours – headbutts, fencing, locomotion. To check classifier accuracy, 30 intervals from a selection of videos for each species were randomly chosen for ground-truthing. A human observer scored each interval for the presence of the behaviour, and the difficulty of identifying that behaviour ('Certain' or 'Difficult'). The table contains the results for each species and behaviour combination. The first number in each cell represents the number of frames scored by the observer and classifier. The second number represents the percentage of frames scored by the manual observer as that behaviour (e.g. for *D. ananassae* headbutts, 32 total frames were scored by the manual observer as 'Behaviour Difficult', where 25 of those were scored as the Behaviour by the classifier and 7 were scored as None by the classifier). Boxes with dark shading indicates agreement between the manual observer and the classifier, while lighter shading indicates disagreement between the observer and classifier (i.e. false negatives in top right of each behaviour: observer scored as behaviour occurring, but the classifier scored as None, and false positives in bottom left: observer scored as None, but the classifier scored as behaviour occurring).

| | | Behaviour | | | | | |
|---|---|---|---|---|---|---|---|
| | | Headbutts | | Fencing | | Locomotion | |
| **Species** | **Human scorer** | Classifier predicted | None predicted | Classifier predicted | None predicted | Classifier predicted | None predicted |
| **ananassae** | Behaviour Certain | 0 (NA) | 0 (NA) | 44 (100%) | 0 (0%) | 32 (87%) | 5 (14%) |
| | Behaviour Difficult | 25 (78%) | 7 (22%) | 62 (83%) | 13 (17%) | 47 (89%) | 6 (11%) |
| | None Certain | 21 (15%) | 122 (85%) | 18 (11%) | 142 (89%) | 62 (30%) | 146 (70%) |
| | None Difficult | 62 (31%) | 140 (69%) | 66 (27%) | 179 (73%) | 66 (31%) | 147 (69%) |
| **bipectinata** | Behaviour Certain | 20 (67%) | 10 (33%) | 41 (68%) | 19 (32%) | 64 (81%) | 15 (19%) |
| | Behaviour Difficult | 30 (68%) | 14 (32%) | 61 (76%) | 19 (24%) | 92 (77%) | 27 (23%) |
| | None Certain | 23 (25%) | 70 (75%) | 34 (21%) | 127 (79%) | 0 (0%) | 127 (100%) |
| | None Difficult | 52 (37%) | 88 (63%) | 61 (29%) | 149 (71%) | 24 (15%) | 133 (85%) |
| **hydei** | Behaviour Certain | 27 (73%) | 10 (27%) | 30 (51%) | 29 (49%) | 180 (84%) | 35 (16%) |
| | Behaviour Difficult | 55 (76%) | 17 (24%) | 42 (44%) | 53 (56%) | 188 (73%) | 69 (27%) |
| | None Certain | 7 (11%) | 59 (89%) | 13 (7%) | 182 (93%) | 11 (4%) | 269 (96%) |
| | None Difficult | 21 (23%) | 69 (77%) | 49 (19%) | 206 (81%) | 11 (3%) | 317 (97%) |
| **nebulosa** | Behaviour Certain | 27 (100%) | 0 (0%) | 18 (100%) | 0 (0%) | 23 (100%) | 0 (0%) |
| | Behaviour Difficult | 36 (100%) | 0 (0%) | 18 (100%) | 0 (0%) | 43 (78%) | 12 (22%) |

| | | | | | | | |
|---|---|---|---|---|---|---|---|
| | None Certain | 35 (12%) | 133 (88%) | 4 (4%) | 89 (96%) | 46 (27%) | 124 (73%) |
| | None Difficult | 39 (29%) | 147 (71%) | 122 (50%) | 124 (50%) | 48 (27%) | 131 (73%) |
| **pseudoobscura** | Behaviour Certain | 58 (77%) | 17 (23%) | 44 (80%) | 11 (20%) | 56 (73%) | 21 (27%) |
| | Behaviour Difficult | 75 (78%) | 21 (22%) | 85 (69%) | 38 (31%) | 62 (60%) | 42 (40%) |
| | None Certain | 22 (21%) | 190 (79%) | 15 (7%) | 188 (93%) | 43 (27%) | 119 (73%) |
| | None Difficult | 33 (21%) | 212 (79%) | 79 (28%) | 207 (72%) | 47 (26%) | 134 (74%) |
| **saltans** | Behaviour Certain | 30 (86%) | 5 (14%) | 10 (100%) | 0 (0%) | 33 (80%) | 8 (20%) |
| | Behaviour Difficult | 68 (81%) | 16 (19%) | 20 (100%) | 0 (0%) | 60 (78%) | 17 (22%) |
| | None Certain | 2 (2%) | 79 (98%) | 9 (6%) | 139 (94%) | 6 (6%) | 99 (94%) |
| | None Difficult | 21 (18%) | 95 (82%) | 87 (32%) | 181 (68%) | 12 (10%) | 105 (90%) |
| **sturtevanti** | Behaviour Certain | 23 (82%) | 5 (18%) | 41 (98%) | 1 (2%) | 77 (85%) | 14 (15%) |
| | Behaviour Difficult | 44 (80%) | 11 (20%) | 61 (98%) | 1 (2%) | 106 (77%) | 32 (23%) |
| | None Certain | 7 (4%) | 164 (96%) | 21 (12%) | 157 (88%) | 4 (2%) | 193 (98%) |
| | None Difficult | 7 (4%) | 183 (96%) | 61 (24%) | 193 (76%) | 8 (4%) | 201 (96%) |
| **willistoni** | Behaviour Certain | 13 (87%) | 2 (13%) | 79 (99%) | 1 (1%) | 35 (100%) | 0 (0%) |
| | Behaviour Difficult | 24 (80%) | 6 (20%) | 156 (86%) | 26 (14%) | 68 (96%) | 3 (4%) |
| | None Certain | 15 (12%) | 113 (88%) | 6 (6%) | 98 (94%) | 24 (18%) | 113 (82%) |
| | None Difficult | 50 (29%) | 124 (71%) | 6 (5%) | 108 (95%) | 24 (18%) | 113 (82%) |

**Table S3: Post-hoc Tukey tests for comparisons of headbutts and fencing between species**
Each cell contains the results of a post-hoc comparison between species (averaged over both mating statuses). The cells contain the odds ratio (calculated as the species in the row/ species in the column), *z ratio*, ***p*-value**. Statistically significant differences are underlined and highlighted in red. Above the diagonal are the comparisons for headbutts and below the diagonal are comparisons for fencing.

| **Species** | ananassae | bipectinata | hydei | nebulosa | pseudoobscura | saltans | sturtevanti | willistoni |
|---|---|---|---|---|---|---|---|---|
| ananassae | | OR: 0.51,z: -*2.09*, **p: 0.42** | 0.26, *-4.578*, **0.0001** | 0.43, *-2.65*, **0.140** | 0.14, *-6.88*, **<0.0001** | 0.23, *-4.91*, **<0.0001** | 0.31, *-3.97*, **0.002** | 1.15, *0.36*, **1.00** |
| bipectinata | OR: 1.23,z: *1.38*, **p: 0.87** | | 0.50, *-3.00*, **0.055** | 0.85, *-0.64*, **1.0** | 0.28, *-6.07*, **<0.0001** | 0.45, *-3.44*, **0.0135** | 0.61, *-2.21*, **0.3478** | 2.27, *2.48*, **0.205** |
| hydei | 0.73, *-2.31*, **0.29** | 0.60, *-3.79*, **0.0038** | | 1.69, *2.41*, **0.24** | 0.55, *-3.62*, **0.0071** | 0.90, *-0.59*, **1.0** | 1.2, *0.99*, **0.977** | 4.49, *4.96*, **<0.0001** |
| nebulosa | 3.43, *-6.16*, **<0.0001** | 2.8, *5.13*, **<0.0001** | 4.69, *8.05*, **<0.0001** | | 0.33, *-5.62*, **<0.0001** | 0.53, *-2.88*, **0.078** | 0.71, *-1.57*, **0.769** | 2.67, *3.04*, **0.0489** |
| pseudoobscura | 1.07, *0.47*, **0.99** | 0.87, *-0.96*, **0.99** | 1.46, *2.90*, **0.071** | 0.31, *-5.93*, **<0.0001** | | 1.63, *2.87*, **0.078** | 2.18, *4.76*, **0.0001** | 8.17, *7.24*, **<0.0001** |
| saltans | 1.42, *2.16*, **0.38** | 1.156, *-0.89*, **0.99** | 1.94, *4.39*, **0.0003** | 0.41, *-4.21*, **0.0007** | 1.32, 1.80, **0.622** | | 1.34, *1.55*, **0.7778** | 5.02, *5.29*, **<0.0001** |
| sturtevanti | 2.11, *-4.55*, **0.0001** | 1.72, 3.3, **0.022** | 2.88, *6.89*, **<0.0001** | 0.61, *-2.29*, **0.297** | 1.97, *4.26*, **0.0005** | 1.49, *2.26*, **0.31** | | 3.74, *4.37*, **0.0003** |
| willistoni | 1.69, *3.20*, **0.03** | 1.38, *1.95*, **0.513** | 2.31, *5.44*, **<0.0001** | 0.49, *-3.32*, **0.0203** | 1.58, *2.87*, **0.079** | 1.20, *1.01*, **0.97** | 0.80, -1.23, **0.92** | |

**Table S4: Post-hoc Tukey tests for comparisons of locomotion between species**

Each cell contains the results of a post-hoc comparison between species (averaged over mating statuses). The cells contain the odds ratio (calculated as the species in the row/ species in the column), *z ratio*, ***p*-value**. Significant differences are underlined and highlighted in red.

| **Species** | bipectinata | hydei | nebulosa | pseudoobscura | saltans | sturtevanti | willistoni |
|---|---|---|---|---|---|---|---|
| ananassae | 1.12, 0.375, **1.0** | 0.1, *-10.41*, **<0.0001** | 1.6, *1.5*, **0.81** | 0.34, -4.65, **0.0001** | 1.08, 0.27, **1.0** | 0.58, *-2.19,* **0.36** | 1.74, 1.76, **0.36** |
| bipectinata | | 0.09, *-9.87*, <**0.0001** | 1.43, 1.08, **0.96** | 0.303, *-4.69,* **0.0001** | 0.966, *-0.112,* **1.0** | 0.52, *-2.44,* **0.22** | 1.56, *1.33*, **0.89** |
| hydei | | | 15.79, 10.82, **<0.0001** | 3.36, *8.38*, <**0.0001** | 10.7, *10.35,* <**0.0001** | 5.75, *10.42,* **<0.0001** | 17.24, *10.99*, <**0.0001** |
| nebulosa | | | | 0.21, *-5.84*, **<0.0001** | 0.68, *-1.22,* **0.93** | 0.36, *-3.62*, **0.0071** | 1.09, *0.26,* **1.0** |
| pseudoobscura | | | | | 3.19, *4.82,* <**0.0001** | 1.71, 2.94, **0.06** | 5.13, *6.08*, <**0.0001** |
| saltans | | | | | | 0.54, *-2.43,* **0.23** | 1.61, *1.48,* **0.82** |
| sturtevanti | | | | | | | 2.99, *3.89*, **0.003** |

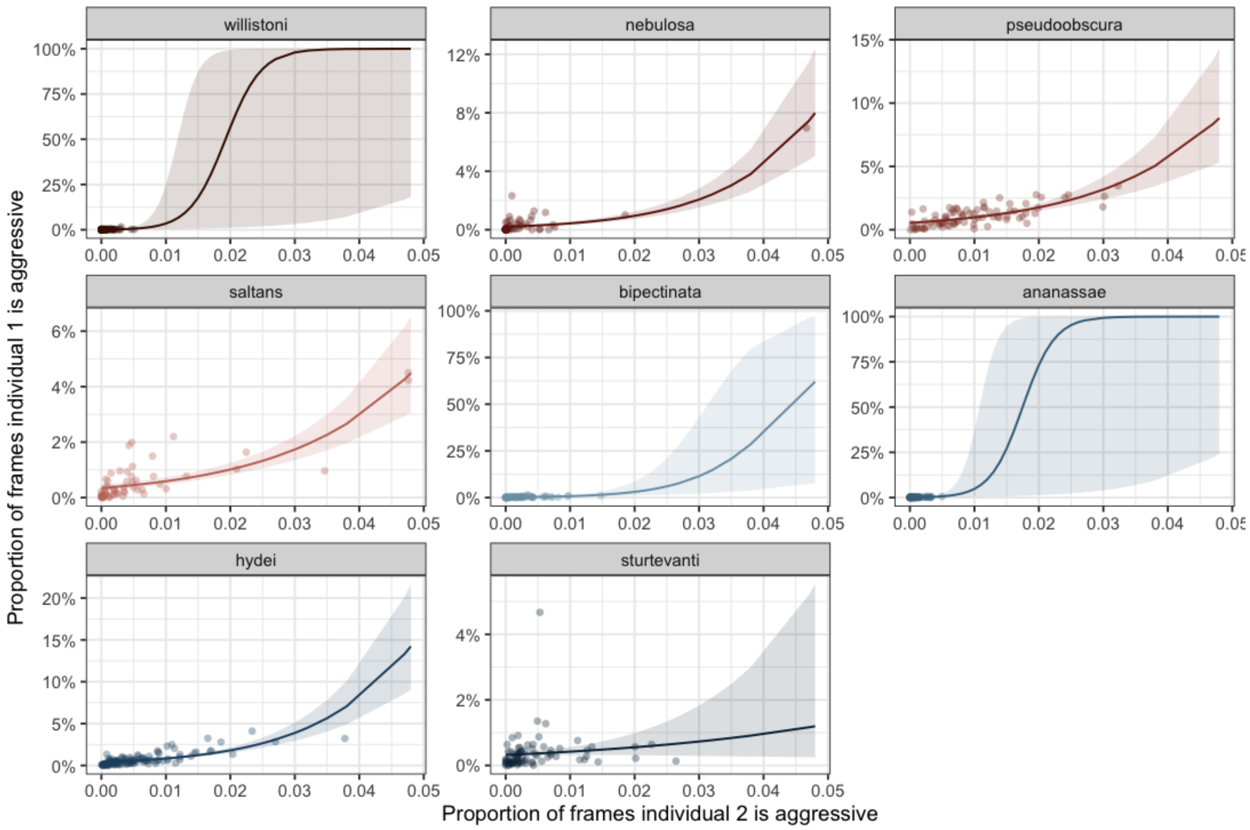


**Figure S1: Correlation of headbutting within pairs by species**
Each point represents a dyad of females with the proportion of frames in a video one individual headbutted on the y-axis and the proportion the other headbutted on the x-axis. Each facet represents data from a different species. Lines represent model predictions, with shaded areas indicating 95% confidence intervals.

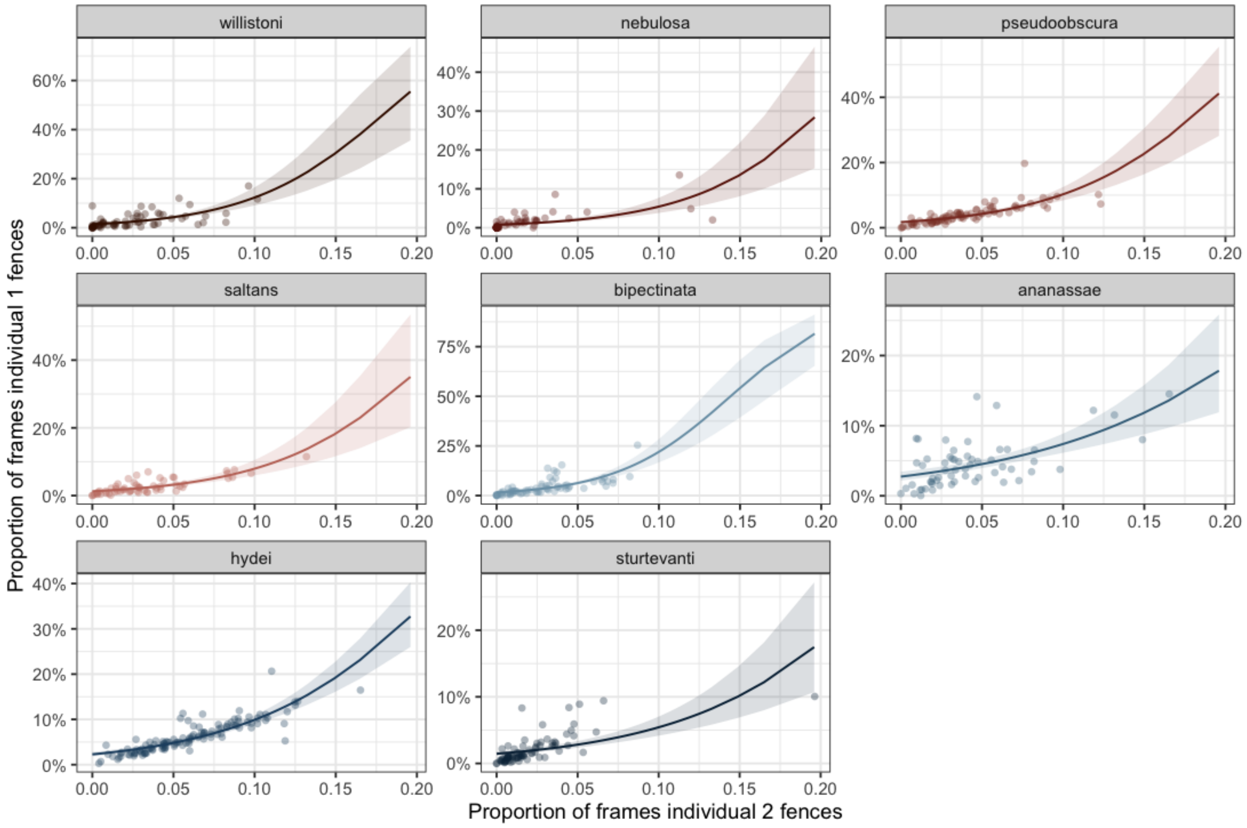


**Figure S2: Correlation of fencing within pairs by species**
Each point represents a dyad of females with the proportion of frames in a video one individual fenced on the y-axis and the proportion the other fenced on the x-axis. Each facet represents data from a different species. Lines represent model predictions, with shaded areas indicating 95% confidence intervals.

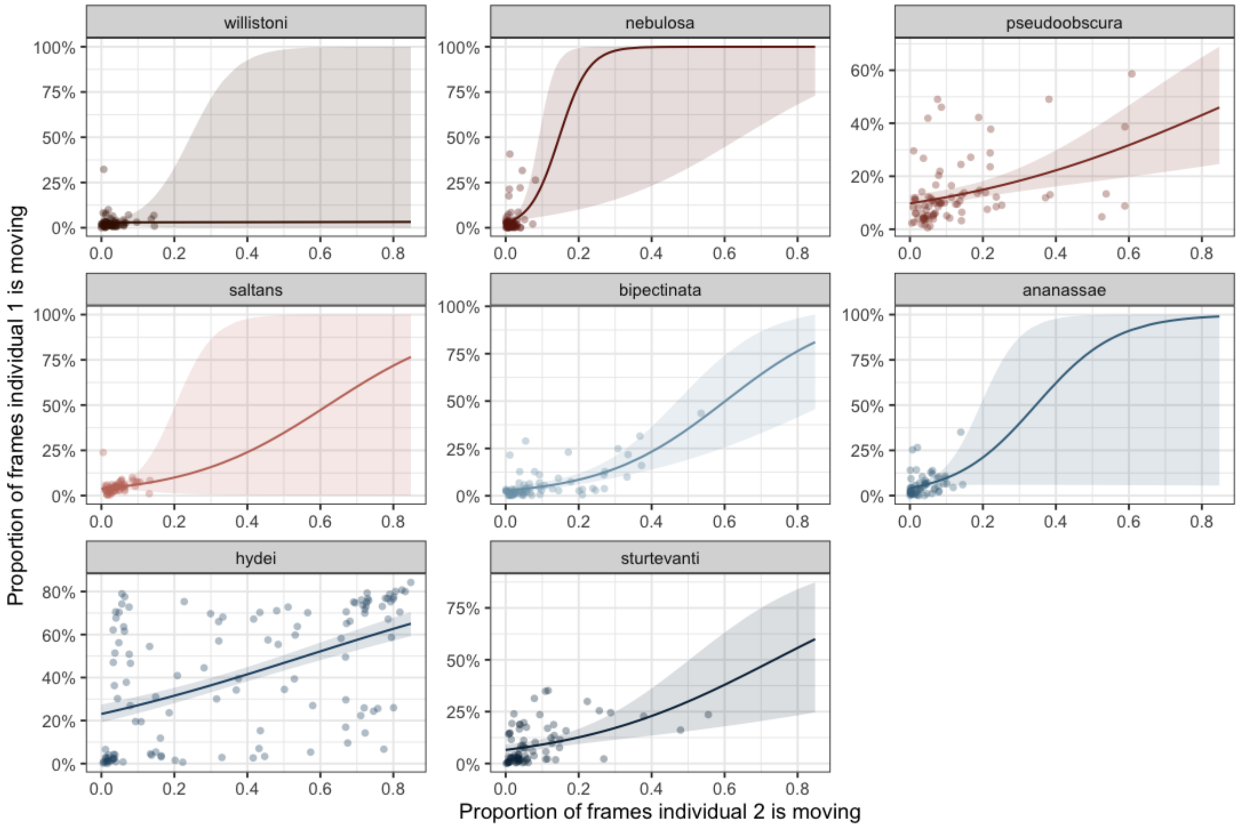


**Figure S3: Correlation of locomotion within pairs by species**
Each point represents a dyad of females with the proportion of frames in a video one individual spent moving on the y-axis and the proportion the other spent moving on the x-axis. Each facet represents data from a different species. Lines represent model predictions, with shaded areas indicating 95% confidence intervals.

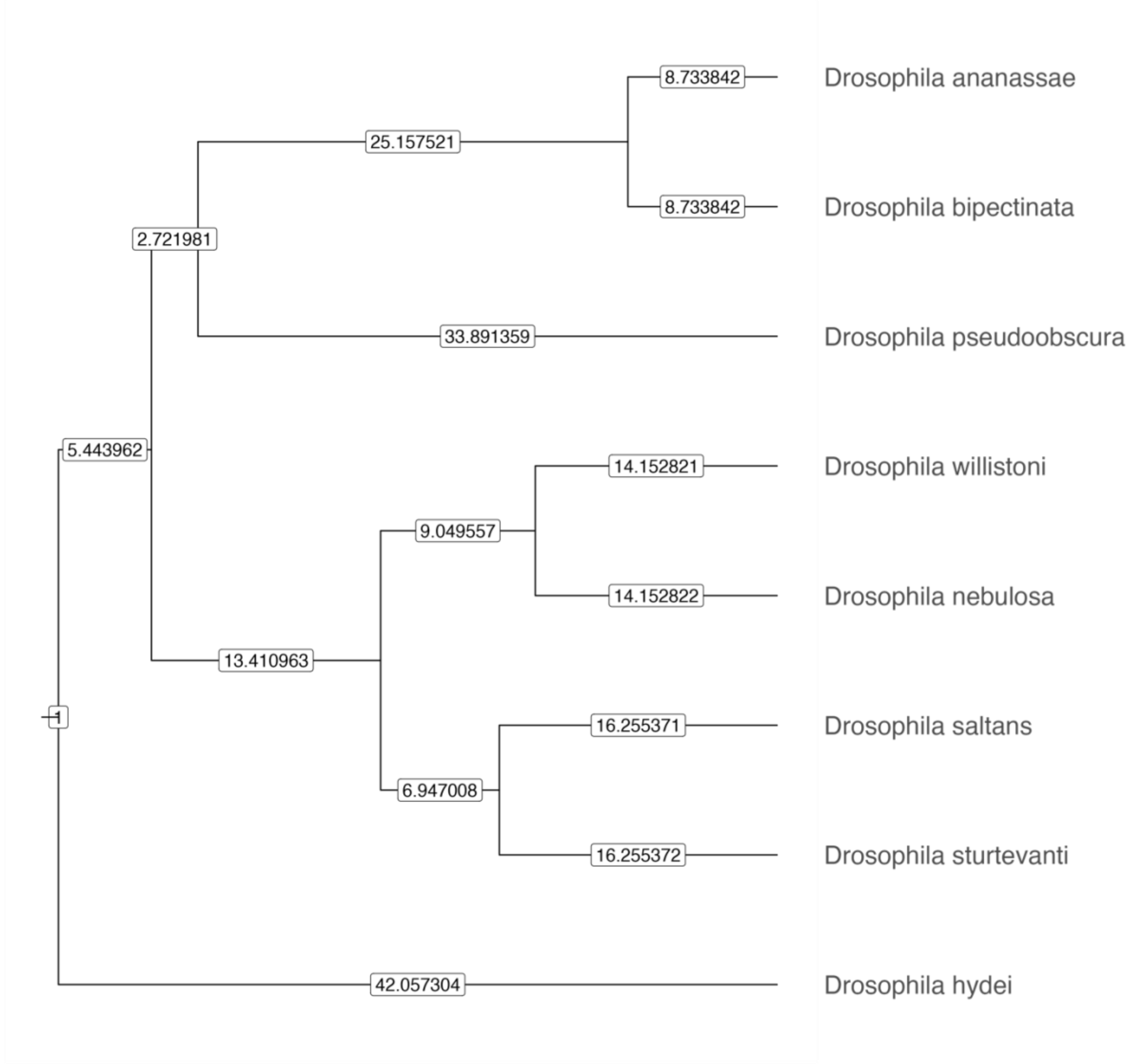


**Figure S4: Phylogeny.**

Phylogeny with branch lengths (in boxes on relevant branches) for the eight species used in our experiment.

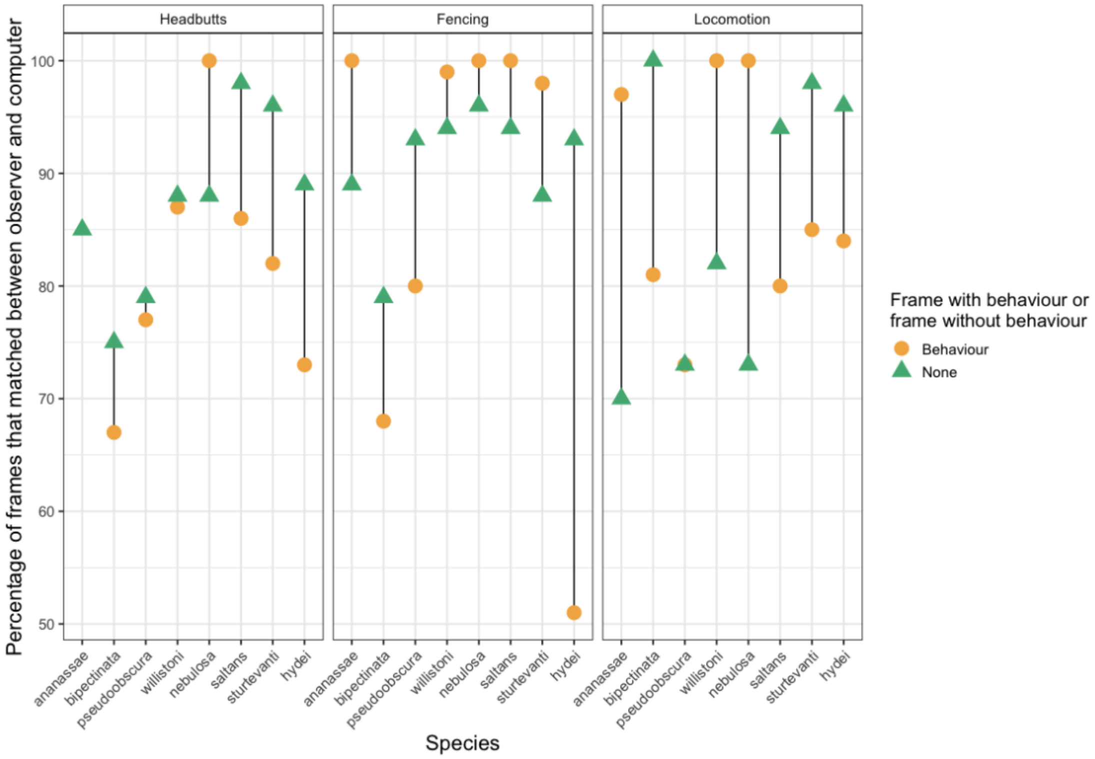


**Figure S5: Machine learning classifier accuracy across species and behaviours.** Yellow circles represent the percentage of frames in the ground-truthing dataset where both the machine learning classifier and the manual observer scored the frame as containing that behaviour. Green triangles represent frames where both the machine learning classifier and the manual observer scored the frame as not containing that behaviour. The lines connect points within a species for ease of visualization across species. These represent the 'Certain' instances of the classifier—those that were judged to be unequivocally containing the behaviour by the manual observer. Results for frames that were less clear are included in table S2. For *Drosophila ananassae*, no frames were scored as containing the 'Certain' headbutting behaviour by either the classifier or the manual observer, so no circle is included. Points higher on the y-axis mean the classifier was more accurate than those lower.

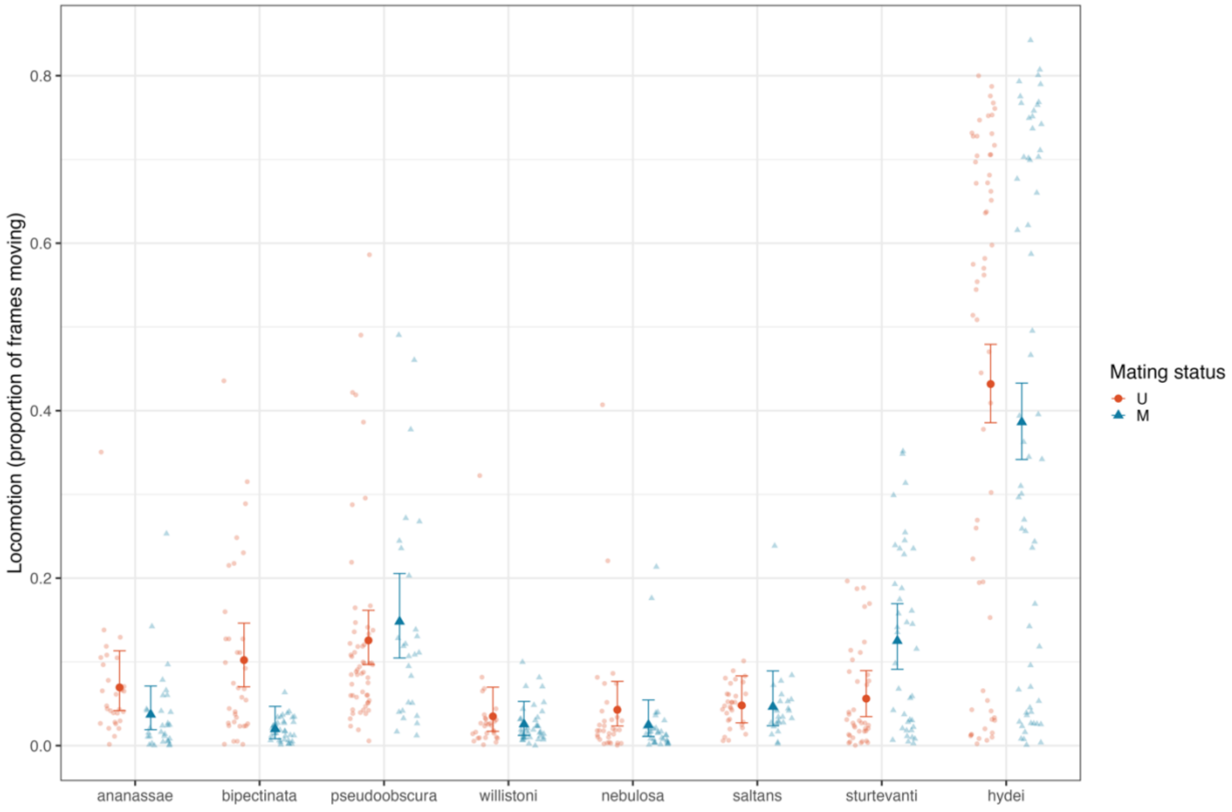


**Figure S6: Proportion of frames spent moving by species and mating status.** Each small point represents one female randomly chosen from a dyad, with orange circles representing unmated pairs and blue triangles representing mated pairs. Large points represent model means with error bars representing 95% confidence interval (asymmetrical due to proportion data).

Species and mating status had a significant interaction in the proportion of frames individual flies were moving ($\chi^2_{7,\,538}$ = 34.569, *p* < 0.0001; Fig. S5). On a species level, *D. hydei* spent significantly more time moving than any other species, around 35% of the 15-minute videos (proportion of frames in video = 0.35 ± 0.02). Within species, the effect of mating status varied significantly. Unmated females moved significantly more than mated females in *D. bipectinata* (OR V/M = 5.64, z = 3.804, *p* = 0.0001), but mated females moved more than unmated females in *D. sturtevanti* (OR = 0.42, z = -3.071, *p*= 0.0021). No other species had a significant difference in locomotion with respect to mating status.

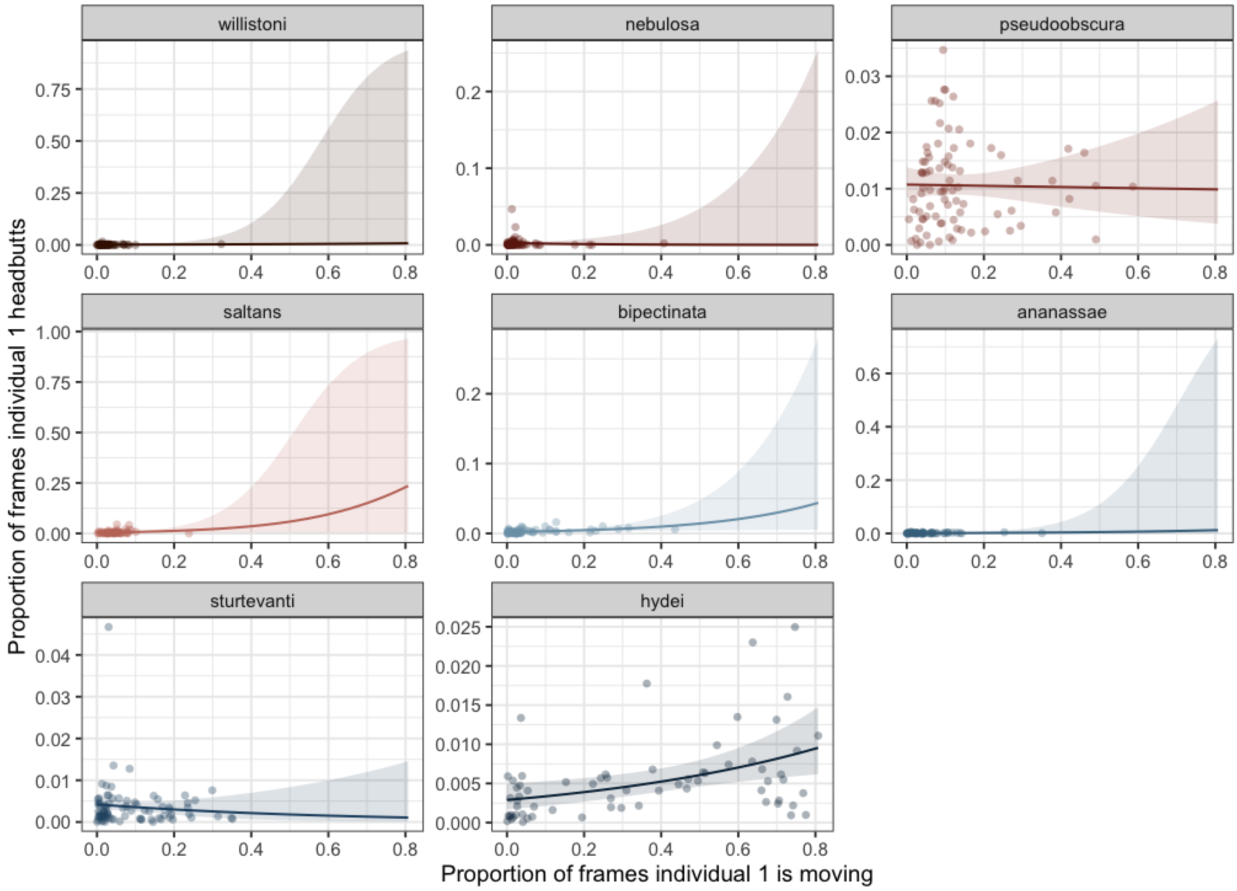


**Figure S7: Relationship between headbutting and locomotion across all species**
Lines indicate model predictions for each species (averaged over mating status). Shaded regions indicate 95% confidence intervals. Each point represents one individual chosen randomly from each dyad. We found no significant effect of locomotion or mating status on headbutts (Locomotion: $\chi^2_{1,\,522}$ = 0.55, *P* = 0.46; Mating status: $\chi^2_{1,\,522}$ = 0.58, *p* = 0.44) and no significant interactions between any of the three variables (Locomotion * Species: $\chi^2_{7,\,522}$ = 4.74, *p* = 0.69; Locomotion * Mating status: $\chi^2_{1,\,522}$ = 0.23, *p* = 0.63; Species * Mating status: $\chi^2_{7,\,522}$ = 12.55, p = 0.08; Locomotion * Species * Mating status: $\chi^2_{7\ 522}$ = 10.71, *p* = 0.15), suggesting a lack of relationship between locomotion and aggression in any of our species or mating status combinations. As in our aggression-only model, we found a significant main effect of species ($\chi^2_{7,\,522}$ = 15.21, *p* = 0.03).

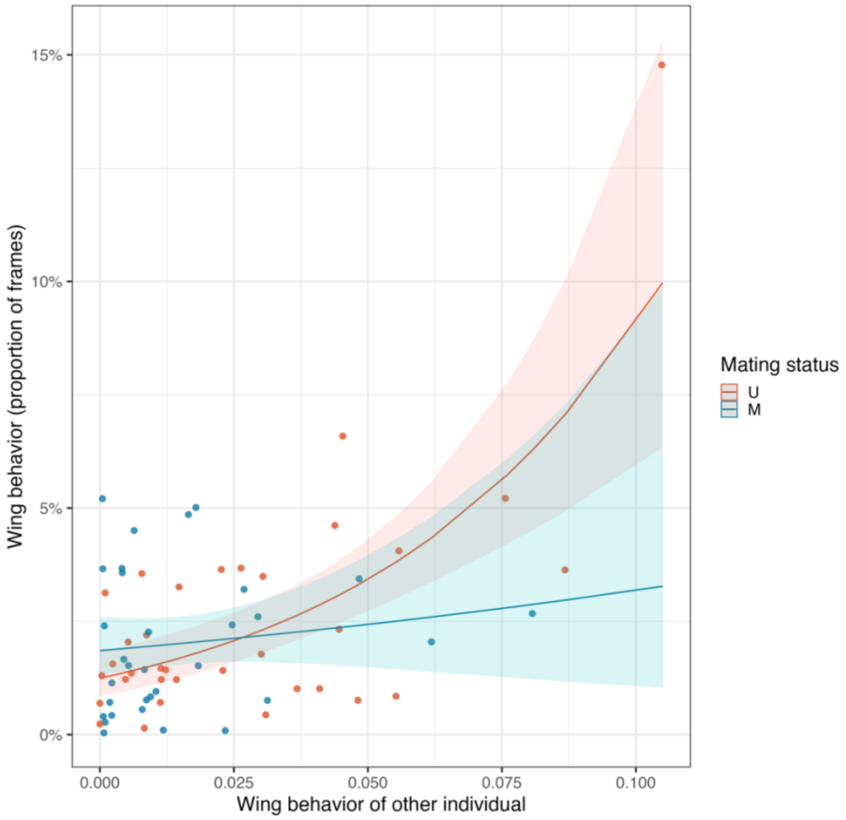


**Figure S8: Correlations within dyads of *Drosophila nebulosa* in wing behaviours**
Lines indicate model results for a generalised linear model with mating status and the proportion of frames the other individual in a pair performed. Intervals indicate 95% confidence intervals. Each data point represents one pair of flies. For unmated females, the number of wing flicks was positively correlated within a pair ($t_{1,62} = 5.72$, $p < 0.0001$), but not in mated females ($t_{1,62} = 0.84$, $p = 0.41$). Unmated females performed the wing behaviour more than mated females ($\chi^2_{1,\,62} = 4.76$, $p = 0.03$).

**Supporting Information**

**Results for fencing**

**Amount of fencing**
We found a significant effect of species on the total proportion of frames in which both flies in a pair were fencing ($\chi^2_{7, 538}$ = 54.92, p < 0.001; fig. S5, table S3). Mating status did not have a significant effect ($\chi^2_{1, 538}$ = 2.58, *p* = 0.11: fig. SI.1), nor did mating status and species interact ($\chi^2_{7, 538}$ = 5.73, *p* = 0.57). The species that spent the most time fencing was *D. hydei* (proportion of frames = 0.11 ± 0.009), while *D. nebulosa* spent the least amount of time (proportion of frames = 0.026 ± 0.004). Closely related species had more similar levels of fencing in unmated females ($\lambda$ = 1.24), while the phylogenetic signal was weaker for mated females, though still present ($\lambda$ = 0.64). However, the relative change in aggression after mating lacked a significant phylogenetic signal ($\lambda$ = - 0.31).

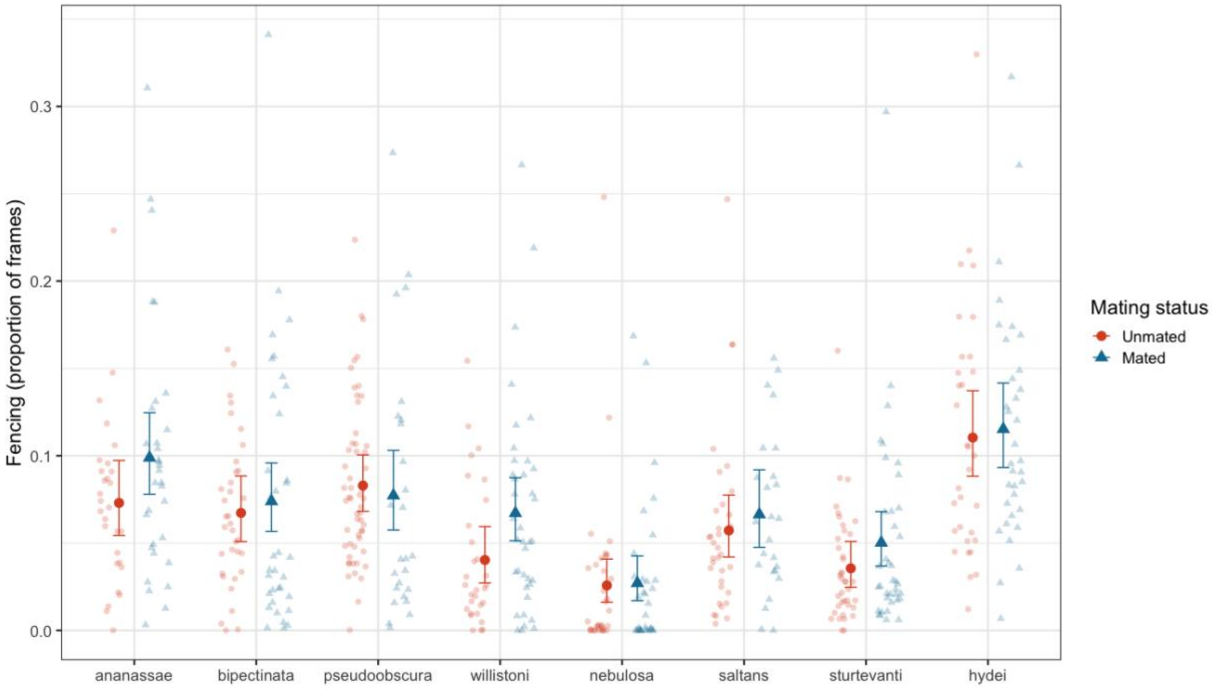


**Figure SI.1: Total proportion of frames scored as fencing for each species and mating status.**
Each small point represents one dyad of females, with black circles representing unmated pairs and gray triangles representing mated pairs. Large points represent model means with error bars representing 95% confidence interval (asymmetrical due to proportion data).

**Fencing and locomotion**
We found no significant effect of locomotion or mating status on fencing (Locomotion: $\chi^2_{1, 522}$ = 0.02, *p* = 0.89; mating status: $\chi^2_{1, 522}$ = 2.59, *p* = 0.11) or significant interactions between any of the three variables (Locomotion * Species: $\chi^2_{7, 522}$ = 13.12, *p* = 0.069; Locomotion * Mating status: $\chi^2_{1, 522}$ = 0.53, p = 0.47; Species * Mating status: $\chi^2_{7, 522}$ = 2.76, *p* = 0.91; Locomotion * Species * Mating status: $\chi^2_{7\ 522}$ = 12.38, *p* = 0.09), suggesting no relationship between locomotion and aggression in any of our species or mating status combinations. As in our aggression-only model, we found a significant main effect of species ($\chi^2_{7, 522}$ = 19.68, *p* = 0.006).

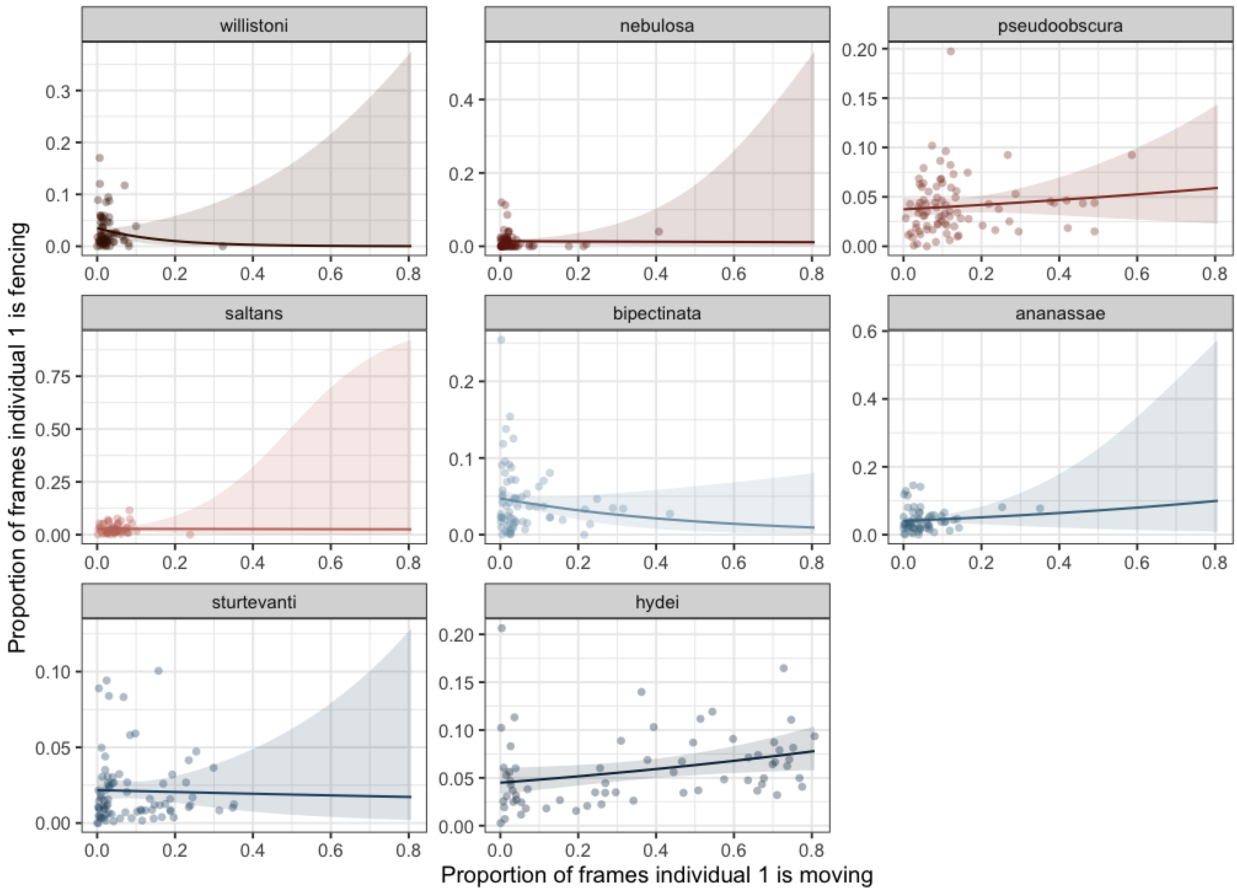


**Figure SI.2: Correlations between fencing and locomotion across all species**
Lines indicate model predictions for each species (averaged over mating statuses). Intervals indicate 95% confidence intervals. Each point represents one individual chosen randomly from each dyad.

**Ovariole number and fencing**

Neither ovariole number nor body size had significant effects on fencing for unmated females (ovariole number: $\chi^2_{1,5} = 0.6$, $p = 0.44$; body size: $\chi^2_{1,5} = 0.17$, $p = 0.68$; $\lambda = 1.14$), or mated females (ovariole number: $\chi^2_{1,5} = 0.2$, $p = 0.66$; body size: $\chi^2_{1,5} = 0.89$, $p = 0.35$; $\lambda = 0.96$). Females in species with fewer ovarioles had higher rates of fencing by mated females. Species with more ovarioles had similar rates of fencing between unmated and mated females, resulting in a negative correlation between ovariole number and mating-induced changes in fencing ($\chi^2_{1,5} = 20.45$, $p < 0.0001$; $\lambda = -0.83$; fig. SI.3). This contrasts with a slight significant positive association between body size and change in fencing after mating ($\chi^2_{1,5} = 7.83$, $p = 0.005$; fig. SI.3).

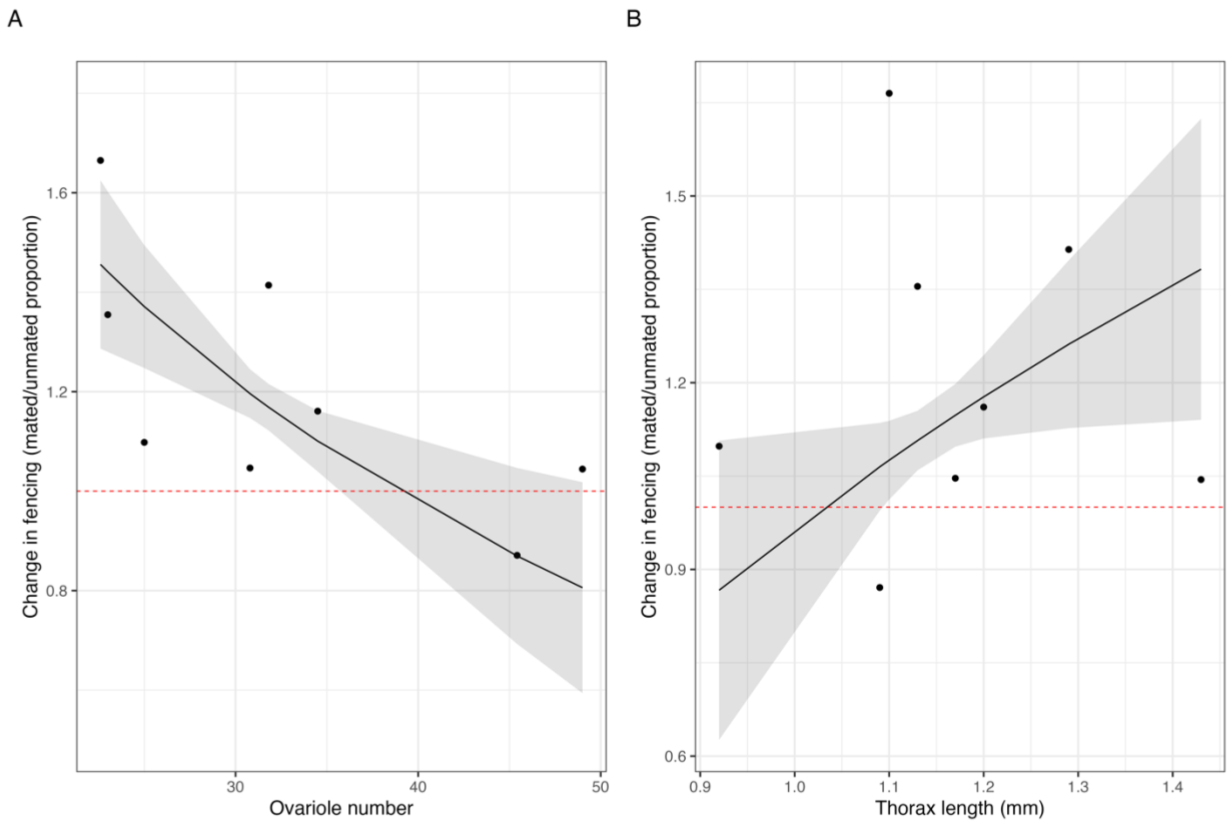


**Figure SI.3: Change in fencing between unmated and mated females is related to ovariole number (A) and thorax length (B) across 8 species**
Change in fencing calculated as mated proportion of frames/unmated proportion of frames. Red dashed line at 1 indicates where mated and unmated females have the same rate of fencing. Points above 1 indicate species where mated females fenced more and points below indicate species where unmated females fenced more. Lines represent model predictions for each variable taking into account the other variable. Intervals are 95% confidence intervals. Neither ovariole number nor body size had significant effects on fencing for unmated females (ovariole number: $\chi^2_{1,5} = 0.6$, $p = 0.44$; body size: $\chi^2_{1,5} = 0.17$, $p = 0.68$; lambda = 1.14), or mated females (ovariole number: $\chi^2_{1,5} = 0.2$, $p = 0.66$; body size: $\chi^2_{1,5} = 0.89$, $p = 0.35$; lambda = 0.96). The difference in fencing between mated and unmated females was associated with ovariole number (A. $\chi^2_{1,5} = 20.45$, $p < 0.0001$; lambda = -0.83). This contrasts with a slight significant positive association between body size and change in fencing after mating (B. $\chi^2_{1,5} = 7.83$, $p = 0.005$).

**Sperm length and fencing**
Sperm length was not associated with unmated or mated female fencing (unmated: $\chi^2_{1,6} = 0.08$, $p = 0.78$; $\lambda = 1.24$; mated: $\chi^2_{1,6} = 0.16$, $p = 0.69$, $\lambda = 0.9$). However, sperm length had a significant effect on the relative change in fencing between mated and unmated females ($\chi^2_{1,6} = 6.08$, $p = 0.01$, $\lambda = -2.52$; fig. SI.4). Females of species with longer sperm fenced more after mating than when unmated.

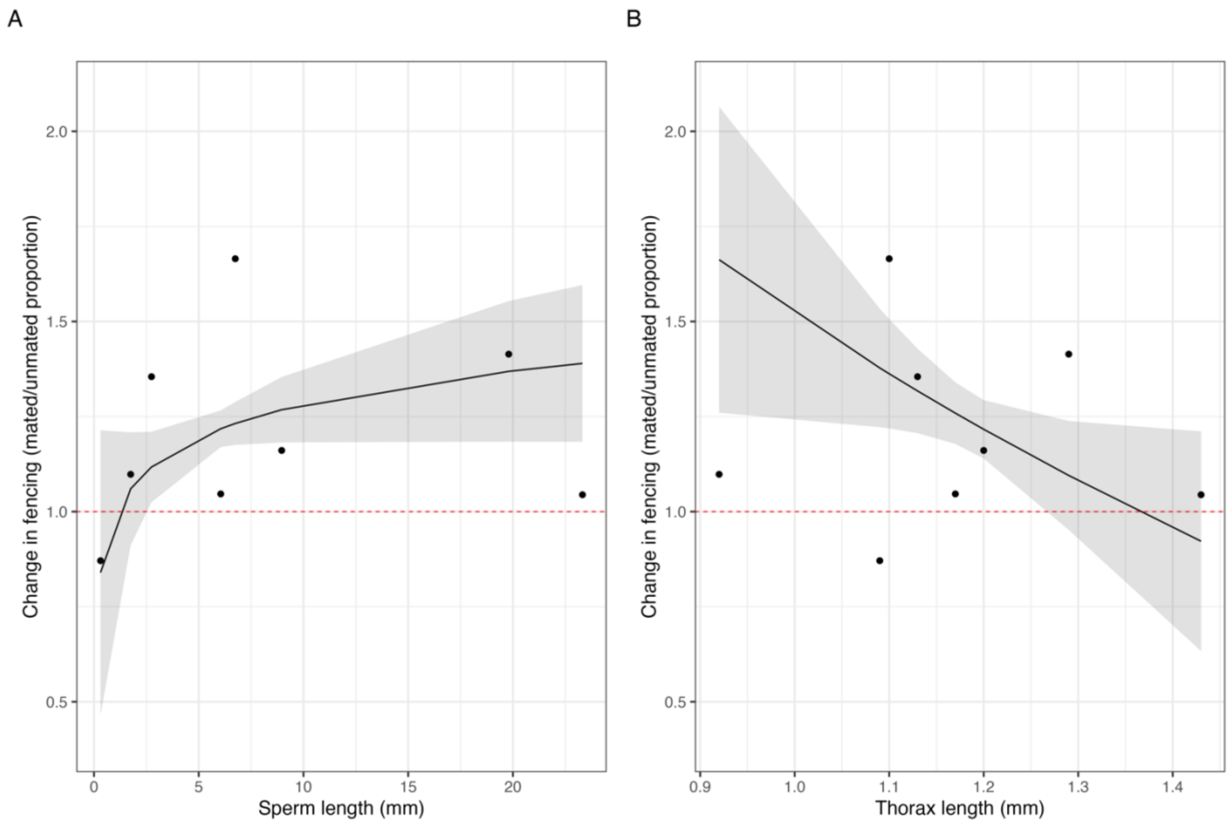


**Figure SI.4: Change in fencing between unmated and mated females influenced by sperm length (A) and body size (B)**
Change in fencing calculated as mated proportion of frames/unmated proportion of frames. The dashed line at 1 indicates where mated and unmated females have the same rate of fencing. Points above 1 indicate species where mated females fenced more and points below indicate species where unmated females fenced more. Lines represent model predictions for each variable taking into account the other variable. Intervals are 95% confidence intervals. Sperm length was not associated with unmated or mated female fencing (unmated: $\chi^2_{1,6} = 0.08$, $p = 0.78$; lambda = 1.24; mated: $\chi^2_{1,6} = 0.16$, $P = 0.69$, lambda = 0.9). However, sperm length had a significant effect on the change in fencing between mated and unmated females (A. $\chi^2_{1,6} = 6.08$, $p = 0.01$, lambda = -2.52;). Females of species with longer sperm fenced more after mating than as unmated females.

**Lifespan and fencing**
There was no relationship between lifespan and mated or unmated fencing (mated: $\chi^2_{1,5} = 0.32$, $P = 0.57$; $\lambda = 0.84$; unmated: $\chi^2_{1,5} = 0.49$, $p = 0.48$; $\lambda = 1.06$).

***Remating rate and aggression***
Remating rate was not correlated with unmated fencing ($\chi^2_{1,5} = 3.17$, $p = 0.08$; $\lambda = 3.61$), mated fencing ($\chi^2_{1,5} = 0.005$, $p = 0.94$; $\lambda = 0.83$), or the relative change in fencing between mated and unmated females ($\chi^2_{1,5} = 1.24$, $p = 0.27$, $\lambda = 0.9$)
